# Integrated metabolic and proteostatic profiling reveals remodeling of proteolytic pathways associated with redox-bioenergetic dysfunction in a PAH^enu2^ mouse model of phenylketonuria

**DOI:** 10.64898/2026.07.08.736353

**Authors:** Francesca Monittola, Elena Perla, Debora Libetti, Antonella Antonelli, Laura Graciotti, Davide Torre, Francesca Pierigé, Anastasia Ricci, Mauro Magnani, Marzia Bianchi, Sara Biagiotti, Luigia Rossi, Michele Menotta, Alessandra Fraternale, Rita Crinelli, Michela Bruschi

## Abstract

Phenylketonuria (PKU) is a genetic metabolic disorder caused by the lack of functional phenylalanine hydroxylase (PAH). Elevated levels of phenylalanine (Phe) are known to be neurotoxic; however, the molecular mechanisms underlying Phe’s effects remain elusive. This study investigates the impact of PKU on proteostasis, redox balance, and metabolism in BTBR PAH^enu2^ mice, a severe disease animal model.

Combined proteomics and metabolomics revealed impaired redox homeostasis in the brain and disrupted mitochondrial energy metabolism (ATP and TCA intermediates). The dysregulation was further supported by decreased levels of ATP, reduced glutathione (GSH), cysteine, and reduced catalase activity. Western blot analyses revealed substantial remodeling of protein degradation systems: the 19S regulatory (Rpt1) subunit and 26S proteasome content and activity were significantly increased, and ubiquitinated protein levels were elevated, indicating protein turnover and activation of the ubiquitin-proteasome system. Autophagy was also activated, as evidenced by a reduced LC3-II/LC3-I ratio, decreased p62 levels, unchanged ATG5 levels, and increased HSPA8 protein expression. By contrast, UPR markers remained stable despite an increase in the oxidized-to-reduced PDI ratio, suggesting a localized shift without activation of a full ER stress response. In parallel, systemic alterations were assessed in whole blood. Indeed, GSH, cysteine, ATP and ADP were decreased in PKU, whereas NADPH increased. These changes were accompanied by reduced activities of GSH reductase and GSH peroxidase, thereby confirming metabolic and redox disruption.

Collectively, these findings indicate that PKU is associated with activation of protein degradation pathways as an adaptive response to cellular stress combined with redox imbalance and energy dysregulation.

**Highlights:**

- Proteasome complex levels and activity were upregulated in PAH^enu2^ mouse brains
- Autophagy was also enhanced in PAH^enu2^ mouse brains
- Despite increased PDI oxidation, the Unfolded Protein Response was not activated
- Redox and energy metabolism were disrupted in both brain and blood

## Introduction

Phenylketonuria (PKU; OMIM*26160) is an inherited metabolic disorder caused by a mutation in the phenylalanine hydroxylase (PAH) gene. PKU results from the enzyme’s inability to convert phenylalanine (Phe) to tyrosine (Tyr), the first and most significant step in Phe catabolism (Bickel et al., 1953). The abnormal accumulation of Phe in untreated PKU infants leads to well-known consequences, the most important of which is substantial disturbance in cognitive development with severe intellectual impairment (Lu et al., 2020). Disruption of amino acid transport and metabolic pathways has been correlated with elevated levels of Phe and/or Phe-related analytes and has been implicated in affecting brain development and function (Embury et al., 2005; Thau-Zuchman et al., 2022). Oxidative damage has been reported in blood cells of patients at diagnosis (untreated phenylketonuric) and in the blood and brain of several mouse models of PKU (Colomé et al., 2003; Sirtori et al., 2005; Sitta et al., 2006). However, little evidence of energy dysregulation was provided for the blood (Ercal et al., 2002). In contrast, in the brain, accumulated Phe negatively affected the activities of catalase (CAT), superoxide dismutase (SOD), and glucose-6-phosphate dehydrogenase (G6PD), decreased the pool of free sulfhydryl groups, and increased the production of reactive oxygen species (ROS) (Bortoluzzi et al., 2019a; Dobrowolski et al., 2022a). In addition, a recent study by Dobrowolski *et al*. reported oxidative stress (OxS) and energy dysregulation in the brain tissue metabolomic profile of PAH^enu2^ mice (Dobrowolski et al., 2022a, 2022b), corroborated by increased ROS and attenuated respiratory chain complex-1 induction in response to pyruvate supplementation, conceivably due to mitochondrial pyruvate transport inhibition.

However, the extent of alterations in biochemical mechanisms associated with PAH deficiency and central nervous system damage remains elusive. Biochemical assessment in experimental and clinical studies involving PKU patients and animal models has primarily focused on amino acids and monoamine neurotransmitters (Dobrowolski et al., 2022a). The proteostasis is critical in nondividing cells, such as neurons, as its failure has been implicated in the development of neurodegenerative diseases [14]. Proteostasis refers to the cellular network that ensures proper protein synthesis, folding, function, trafficking, and degradation. Amino acid deprivation, such as the Phe restriction occurring in PKU, could impair translation and protein folding, leading to proteotoxic stress (Costa-Mattioli and Walter, 2020). The Unfolded Protein Response (UPR) is an adaptive mechanism activated in the ER upon accumulation of misfolded proteins, involving three branches— protein kinase-like ER kinase (PERK), inositol-requiring kinase 1α (IRE1α), and activating transcription factor 6 (ATF6). Key markers include phosphorylated eIF2α and ATF4 (PERK pathway), spliced XBP1 (IRE1α pathway), and cleaved ATF6. In ER-unstressed cells, the abundant ER-resident chaperone BiP binds to the luminal domains of all three receptors, keeping them inactive [38,39]. These pathways reduce global protein synthesis, upregulate ER chaperones, and enhance degradative processes to restore proteostasis (Bobak et al., 2016). Moreover, PERK could activate ATF4 by inhibiting heat shock protein 90 (Hsp90), leading to IRE1α degradation and dephosphorylation of eIF2α, thereby driving the cytoprotective UPR into an apoptotic pathway [41].

The removal of misfolded and aggregated proteins is accomplished by two major catabolic routes: the ubiquitin-proteasome (UPS) and the autophagy-lysosomal systems. The proteasome is a protein complex consisting of a 20S core, a proteolytic cylinder composed of two α-rings and two β-rings. Among the β-rings, the β1, β2, and β5 subunits respectively exhibit caspase-like, trypsin-like, and chymotrypsin-like activities. A specialized proteasome variant arises in response to pro-inflammatory stimuli, in which the constitutive catalytic β subunits are replaced by the inducible β1i, β2i, and β5i subunits, forming the immunoproteasome (i20S) (Monittola et al., 2025). Both 20S cores can be capped by regulatory particles that modulate their activity. When capped by a 19S particle, they form the canonical 26S proteasome (when capped by two 19S particles, they form a 30S proteasome), a complex that is ATP- and ubiquitin (Ub)-dependent. In this case, proteins destined for proteasomal degradation are first tagged with Ub molecules. Another regulatory complex is PA28αβ, which allows degradation in an ATP- and Ub-independent manner (Tanaka, 2009). On the other hand, during autophagy, cytoplasmic components are sequestered into a double-membrane vesicle called an autophagosome, which fuses with lysosomes to form autolysosomes, where the cargo is degraded. Evaluation of autophagic flux relies on key molecular markers: LC3-I is lipidated to LC3-II during autophagosome formation, and the LC3-II/I ratio reflects the degradation rate; ATG5, as part of the ATG12–ATG5–ATG16L1 conjugation system, is required for LC3 lipidation and autophagosome elongation. Moreover, autophagy and proteasomal degradation are linked through the selective receptor p62/SQSTM1, which binds ubiquitinated cargo and LC3-II, and directs them for autophagic degradation. It is degraded within autolysosomes; thus, its accumulation indicates impaired autophagy (Ortega et al., 2024).

Proteostasis and redox dysregulation are interconnected processes. Indeed, both catabolic systems may be modulated by Nrf2-mediated redox control, and the accumulation of damaged proteins can also exacerbate OxS (Buttari et al., 2025). Whether neuronal dysfunction in PKU is accompanied by altered proteostasis remains to be investigated. In the present study, we aimed to evaluate the concomitance of metabolic alterations, as reflected in metabolomic and proteomic profiles, with disturbances of proteostasis in the brain of PAH^enu2^ mice, along with enzymatic and non-enzymatic antioxidant defense parameters in both blood and brain.

For this purpose, a mouse model harboring nucleotide substitutions in the murine PAH gene was exploited. The loss-of-function variant of the murine PAH enzyme c.835T>C (p.Phe263Ser) results in a severe classical PKU phenotype in PAH^enu2^ mice that closely resembles untreated human PKU (Eichinger et al., 2018). Thus, a better understanding of proteostasis changes and redox imbalance in brain tissues in PKU may help elucidate the molecular mechanisms underlying neurological dysfunctions or damage, providing a base for future studies aimed at identifying new therapeutic targets.

## Materials and Methods

### Animal model

BTBR mice were raised in the animal shelter of the Biochemistry and Biotechnology section of the Department of Biomolecular Sciences at the University of Urbino Carlo Bo. PAH^enu2^ (PKU ^−/−^) and wild type (WT ^+/+^) mice used in this study were obtained by mating heterozygous animals from the BTBR strain. Animals were housed in standard cages (4 mice per cage) under a 12-h light-dark cycle and controlled conditions of temperature (22 ± 1 °C), humidity (60%), and air changes (every 12 h). Genetic characterization was performed as previously described (McDonald and Charlton, 1997). All mice were fed a Teklad Global 18% Protein Rodent Diet (Teklad, Harlan Laboratories Inc., Madison, WI) and had water *ad libitum*. for the protection of animals used for scientific purposes. All experiments were conducted in accordance with the Guidance on the operation of the Animals Act 1986 and associated guidelines, EU Directive 2010/63, Italian national legislation (DL26/2014) governing the use of animals for research, and the guidelines of the National Institute of Health on the use and care of laboratory animals (Authorization n° 486/2017-PR). All surgical procedures were performed under inhaled isoflurane general anesthesia. Animals were sacrificed by rapid cervical dislocation at 6 months of age (weight between 30 – 40 g). Blood was collected by terminal bleeding, and plasma was prepared by centrifugation. Brain tissue was dissected, snap-frozen in liquid nitrogen, and half-brains were pulverized under liquid nitrogen to a fine, homogeneous powder.

### Metabolomics and proteomics

Pulverized half-brains (WT, n = 6, PKU n = 6) were processed using ice-cold PBS and harvested in cold 80/10/10 LC/MS grade methanol/acetonitrile/water (Carl Roth, Karlsruhe, Germany). Insoluble material was pelleted by centrifugation at 20,000 g for 20 min. An acidic extraction buffer containing 20 mM ammonium acetate and 2.0 mM Na-ascorbate in 100% methanol was prepared to preserve redox-sensitive metabolites in parallel with global metabolic profiling of mammalian cells, following the protocol established by Petrova and colleagues (Petrova et al., 2021).

Samples were analyzed using an ultra-high-pressure liquid chromatography Vanquish system (Thermo Fisher Scientific, USA) coupled to a mass spectrometry Exploris 240 (Thermo Fisher Scientific) following previously published instructions (Monittola et al., 2025).

Compounds were separated by both C18 Hypersyl GOLD column (150×2.1mm x 1.9µm) and by the amide-HILIC Column (150×2.1mm 2.6 µm both from Thermo Fisher Scientific). Acquisitions were performed in both positive- and negative-ion polarity modes. The Exploris 240 was set to the MS1 range of 80-900 m/z, 120,000 resolution at m/z 200, AGC target of 10e6, and auto maximum injection time. For MS2, auto m/z range, stepped HCD normalized collision energy (20, 50, 80, 150%), and 30,000 resolution at m/z 200, AGC target 2e5, and maximum injection time 70ms. Calibration was performed before each analysis sequence, and the internal calibrant was used in run-start mode. Untargeted metabolomics was performed using the deep scan AcquireX software, with 5 ID runs, 5 quality controls, and 3 replicates per sample for statistical analysis. Raw data were processed by Compound Discoverer software Ver 3.3 (Thermo Fisher Scientific).

Protein pellets were dissolved in the EasyPep lysis buffer (Thermo Fisher Scientific, USA) and 50 µg were subsequently processed by the EasyPep™ Mini MS Sample Kit (Thermo Fisher Scientific). Purified peptides were employed in label-free bottom-up proteomic analysis. Briefly, two µg of peptides from each experimental condition were injected into the UltiMate™ 3000 RSLC nanoHPLC coupled to a second Orbitrap Exploris 240 Mass Spectrometer. Peptides were inline desalted by Acclaim PepMap C18 trap Column (5 μm, 0.3 mm x 5 mm Thermo Scientific), and then resolved by the Easy Spray Pepmap RSLC C18 (2 μm, 75 μm X 50Cm) at a flow rate of 250nl/min by a linear gradient of phase B (80% acetonitrile/0.1% formic acid, solvent A was 0.1% formic acid in water) from 2% to 50% in 200 min, to 95% in 20 min, kept for 10 min and then the column was re-equilibrated for 10 min. Data were acquired in a positive mode and in a data-dependent manner. For MS1 m/z range was set to 350-1500 at 120,000 resolution (at m/z 200), AGC target 3e6, and auto maximum injection time. For MS2, switch to DDA when the ion intensity was above 1e4, with a m/z range in auto mode. The top 20 strategy was adopted with a dynamic exclusion time of 80 seconds and a 10 ppm tolerance. HCD normalized collision energy was set to 30%, AGC target 1e5, and maximum injection time 40ms. Resolution was set to 15,000 at m/z 200, and an internal calibrant was employed in run start mode. Raw data were analyzed using Proteome Discoverer software v2.5 (Thermo Fisher Scientific, USA), employing a label-free signal quantification strategy.

### Computational metabolomics and proteomics data analysis

Processed metabolomics and label-free proteomics output tables were further analyzed using a custom Python workflow to enable integrated comparison of PKU and WT samples. The workflow imported quantitative metabolite and protein tables, harmonized column names, standardized statistical fields, and generated curated output files for downstream single-omics and multi-omics analyses. For each molecular feature, log2 fold changes, false discovery rate (FDR) adjusted values, and −log10 p values were computed. Differentially regulated metabolites and proteins were classified as increased or decreased in PKU relative to WT using the predefined statistical thresholds for the study, including an FDR < 0.05 and fold-change-based filtering.

Metabolites and proteins were first analyzed independently and assigned to broad functional families or biological modules to obtain a higher-level representation of pathway-level remodeling. For each metabolite or protein family, the workflow calculated the number of detected features, the number of nominally significant features, the mean and median log2 fold change, and a directional z-like activity score based on the distribution of log2 fold changes within each omics layer. This single-layer module score was used to summarize the direction and internal consistency of metabolite or protein changes within each functional category.

In addition to single omics summaries, a dedicated multi-omics integration step was performed to identify coordinated alterations across the metabolome and proteome. Metabolite and protein changes were mapped to shared biological modules and to enzymatic reactions. The integrated workflow generated reaction- and module-level activity tables, combining metabolomic and proteomic evidence to infer the direction of pathway remodeling in PKU relative to WT.

Directional activity z-scores were then calculated from the integrated multi-omics output to prioritize reactions or functional modules showing coherent evidence across molecular layers. Positive activity z-scores indicated reactions or pathways globally increased in PKU relative to WT, whereas negative scores indicated globally decreased activity. For each activity state, the workflow retained the supporting molecular evidence, allowing the distinction between concordant metabolome–proteome changes and weaker or heterogeneous signals. These integrated z-score tables were used to identify coordinated pathway-level alterations. UniProt was used to retrieve gene symbols, protein names, and enzyme commission annotations; KEGG REST was used to map metabolites and proteins to metabolic reactions; g:Profiler was used for functional enrichment of protein lists, and STRING was used to generate protein interaction information. These annotations were used to explore PKU-associated molecular alterations at the level of functional modules, enzymatic reactions, enriched pathways, and interaction networks.

### HPLC-based quantification of brain thiols and ATP

Brain cortexes (∼ 20-30 mg) (WT, n = 6 and PKU, n = 6) were swiftly set in 300 μL of ice-cold precipitating solution (100 mL containing 1.67 g of glacial metaphosphoric acid, 0.2 g of disodium EDTA, and 30 g of NaCl) before being homogenized for 30 sec using an IKA T10 Ultra-Turrax^®^ homogenizer (IKA-Werke GmbH, Germany) and sonicated for 1 min at 50 W.

The sample was kept on ice for 10 min, then centrifuged at 13,800 × g for 10 min at 4°C. Supernatant was frozen and stored at −20°C, while the pellet was discarded. Before HPLC injection, 15 μL Na_2_HPO_4_ (0.3 M) and 7.5 μL 5,5′-dithiobis (2-nitrobenzoic acid) (DTNB), known as Ellman’s reagent, were added to 60 μL of the supernatant. DTNB solution was prepared by dissolving 20 mg of DTNB in 100 mL of sodium citrate solution (1% w/v). The mixture was used for cysteine and GSH determination by HPLC on a Vanquish system (Thermo Scientific, USA) at a wavelength of 330 nm. Separation and elution conditions were previously described elsewhere (Bruschi et al., 2025; Masini et al., 2025).

For ATP determination, a BDS Hypersyl TM C18 column (250 mm x 4.6 mm, particle size 5 μm) (Thermo Scientific, USA) was used, coupled with a Teknokroma Tracer Excel ODS guard column (Teknokroma Analitica SA, Spain). The mobile phase consisted of buffer A (0.1 M KH_2_PO_4_, pH 6.0) and buffer B (10% v/v methanol in buffer A). The elution program was as follows: 100% buffer A for 9 minutes, 75% buffer A from 9 to 15 minutes, 10% buffer A from 15 to 17.5 minutes, 100% buffer B from 17.5 to 19.5 minutes, kept for an additional 6 minutes before switching back to 100% buffer A from 25.5 to 30 minutes, and continued up to 36 minutes. The injection volume was 50 μL, with a flow rate of 1.3 mL/min. Detection was performed at 254 nm using a Jasco HPLC LC-net II (Jasco Inc., Japan). Quantitative measurements were compared with known concentration standards and normalized to the specimen weight.

### Gene expression (RT-qPCR)

WT and PKU brain lysates from 180 PND (WT n=6, PKU n=6) were prepared in Buffer RLT Plus, and total RNA from each sample was extracted using the RNeasy Plus Mini Kit (QIAGEN) according to the manufacturer’s instructions. RNA (500 ng) was reverse-transcribed using PrimeScript™ RT Master Mix (Perfect Real Time; Takara Bio Europe SAS). cDNA was amplified with PowerTrack™ SYBR Green Master Mix (Applied Biosystems™) and visualized with SYBR Green on a QuantStudio™ 5 Real-Time PCR System (Applied Biosystems, Thermo Scientific, USA). Thermal cycling was performed as follows: 2 min at 95 °C; 40 cycles of denaturation at 95 °C for 15 s, annealing at 60 °C for 1 min, and extension at 60 °C for 1 min. Expression data were calculated using the 2^-^ΔΔCt^ method (Livak and Schmittgen, 2001).

### Western immunoblotting analysis

The brains (WT, n = 6; PKU, n = 6) were pulverized, frozen with liquid nitrogen, and the powder was immediately homogenized in specific lysis buffers for subsequent analysis. For SDS-PAGE, the denaturing buffer consisted of 50 mM Tris–HCl, pH 7.8, 0.25 M sucrose, 2% (w/v) sodium dodecyl sulfate (SDS), 10 mM N-ethylmaleimide, supplemented with a cocktail of protease (Complete, Roche) and phosphatase inhibitors (1 mM NaF, 1 mM Na_3_VO_4_). Samples were heated at 100 °C, sonicated at 70 W for 40 s, and centrifuged at 14,000 × g to remove debris. Proteins were separated by SDS PAGE (polyacrylamide gel electrophoresis) and immunoblotted onto PVDF (polyvinylidene difluoride) 0.2 µm pore size membrane. After transfer, the total amount of loaded protein was detected using the No-Stain™ Protein Labeling Reagent (#A44717, Invitrogen) following the manufacturer’s instructions and visualized with the ChemiDoc MP system (Bio-Rad) using Cy3 acquisition settings (Green epi 602/50 filter). After blocking, membranes were incubated with the following primary antibodies: proteasome 20S α1, 2, 3, 5, 6 and 7 subunits (pan α) (MCP231) (#PW8195, Enzo Life Sciences), PSMB5/β5 (#ALS17241, Abcepta), PSMB8/β5i (#13635, Cell Signaling Technology), PSMB7/β2 (#14771, ABclonal), PSMC2/ATPase 2 (#A1985, ABclonal), ubiquitin (kindly provided by Prof. A.L. Haas, New Orleans School of Medicine); PSMB6/ β1 (**#**13267, Cell Signaling Technology); LC3B (#2775, Cell Signaling Technology); p62/SQSTM1 (#5114, Cell Signaling Technology); HSPA8 (#8444, Cell Signaling Technology); ATG5 (A0203, ABclonal); (#5536, Cell Signaling Technology); Phospho-mTOR (Ser2448) (#5536, Cell Signaling Technology); mTOR (#2983, Cell Signaling Technology); UBE1 a/b (#4891, Cell Signaling Technology); Phospho-eIF2 alpha (Ser51) (#3398, Cell Signaling Technology); eIF2 alpha (#5324, Cell Signaling Technology); BiP/grp78 (#3177, Cell Signaling Technology); PDI (#3501, Cell Signaling Technology); Phospho-IRE1-S724 (#AP0878, ABclonal); IRE1 alpha (#3294; Cell Signaling Technology); ATF-6 (#65880, Cell Signaling Technology); Hsp90α/β (#A5027, ABclonal). Immunoreactive bands were detected using a horseradish peroxidase-conjugated secondary antibody (Bio-Rad Laboratories Inc.) and the WesternBright ECL enhanced chemiluminescence detection kit (Advansta) on a ChemiDoc MP Imaging System (Bio-Rad). The immunoreactive bands were quantified using the Image Lab analysis software version 5.2.1 (Bio-Rad). Raw data and loading controls are provided in the Supplementary material (S5 – S6).

### Protein carbonylation assay

Powdered brain tissue obtained from WT and PKU mice was homogenized in a lysis buffer containing 10 mM HEPES/KOH (pH 7.9), 10 mM KCl, 0.2 mM ethylenediaminetetraacetic acid (EDTA), 0.01% (v/v) Nonidet P-40, and 50 mM dithiothreitol (DTT), supplemented with a protease inhibitor cocktail. Protein carbonylation was assessed using the OxyBlot™ protein oxidation detection kit (Sigma‒Aldrich, #S7150) according to the manufacturer’s protocol. Briefly, 5 μg of each sample was treated with 1× 2,4-dinitrophenylhydrazine (DNPH) solution, while negative control samples were incubated with neutralization solution. Proteins were separated on a 12% (w/v) acrylamide gel and subsequently transferred onto a membrane for Western blot analysis using the antibodies included in the kit.

### Native PAGE analysis of proteasome complexes

Native gel analysis of proteasome complexes was performed as described in Yazgili *et al*. (2021) and Roelofs *et al*. (2018) (Roelofs et al., 2018; Yazgili et al., 2021). Powdered brains, previously described, were homogenized under native conditions in a lysis buffer consisting of 10 mM Tris–HCl, pH 7.5, 5 mM MgCl_2_, 10 mM NaCl, 10% (v/v) glycerol, 1 mM dithiothreitol (DTT), 2 mM ATP, and a cocktail of protease inhibitors. Lysates were centrifuged at 20,000 × g at 4 °C. 15–20 µg of protein was loaded onto 4% (w/v) polyacrylamide gels in Tris/Borate buffer containing 0.5 mM ATP. Gels were run at 150 V for 2.5 h at + 4 °C. In gel, proteasome activity was performed by incubating the gel in reaction buffer (50 mM Tris–HCl, pH 7.5, 1 mM ATP, 10 mM MgCl_2_, 1 mM DTT, 50 μM fluorogenic substrate) for 30 min at 37 °C in the dark. Images were visualized in a Gel Doc system (Bio-Rad) under UV light. After denaturation in solubilization buffer (2% SDS, 66 mM Na_2_CO_3_, 1.5% β-mercaptoethanol) for 20 min at room temperature, proteins were blotted onto PVDF membranes and stained with antibodies against proteasome subunits.

### Proteasome activity assay

Proteasome activity was measured in native cell extracts obtained as described above using the following synthetic fluorogenic substrates (Cayman Chemicals): s-LLVY-AMC (Suc-Leu-Leu-Val-Tyr-7-amido-4-methylcoumarin) for the chymotrypsin-like activity; Boc-LRR-AMC (Boc-Leu-Arg-Arg-AMC) for the trypsin-like activity, and Z-LLE-AMC (Z-Leu-Leu-Glu-AMC) for the caspase-like activity. The assay buffer consisted of 50 mM Hepes/KOH (pH 7.8), 10 mM KCl, 2.5 mM ATP, and 25 mM MgCl_2_. The reaction was initiated by the addition of the fluorogenic peptide (200 μM s-LLVY-AMC, Z-LLE-AMC, and Boc-LRR-AMC). Protein extract concentration was within the linear signal-concentration range, typically between 0.2 and 0.1 μg/μL. After hydrolysis, the release of AMC from peptidyl derivatives was measured at 37 °C for 30 min with excitation/emission wavelengths of 355/460 nm. Proteasome activity was calculated from the slope after linear regression analysis of the values plotted as a function of time (R_2_ > 0.98).

### Redox enzyme activities

Glucose-6-phosphate dehydrogenase (G6PD), superoxide dismutase (SOD), and catalase (CAT) activities were quantified from pulverized brain sections (PKU n=6, WT n=6) using the respective Elabscience Assay Kits (E-BC-K056-M, E-BC-K019-M, E-BC-K031-M, respectively) according to the manufacturer’s instructions. Enzyme activity was normalized on total protein content (measured using the Bradford Assay) and was expressed as Units/mg of protein.

For the enzyme assessment of G6PD, glutathione peroxidase (GPX) activity, and glutathione reductase (GR) in blood samples from 6 PKU and 6 WT, hemolysates were prepared by adding 2 volumes of a stabilizing solution (pH 7.4; phosphate buffer, 3 mM; mercaptoethanol, 3 mM; EDTA, 0.5 mM) and incubating on ice for 1 h. Hemolysate samples were processed according to Beutler’s procedure (Beutler E., 1984). Quantification was based on Drabkin’s Assay for normalization on hemoglobin content and expressed as Units/mg of hemoglobin.

### Histological Evaluation

Mouse brains were carefully removed after euthanasia and immediately immersed in 10% neutral buffered formalin. Following fixation, tissues were dehydrated through a graded ethanol series, cleared in xylene, and then paraffin infiltrated and embedded. Blocks were trimmed and sectioned using a rotary microtome. Coronal and horizontal sections of the cortex and cerebellum were cut at 4-5 μm thickness. Slides were deparaffinized and rehydrated, then stained with hematoxylin and eosin (H&E) to visualize cytoplasm and cell nuclei, enabling detailed interpretation of cellularity and the neuronal network. H&E-stained sections were examined using a Nikon Eclipse 80i microscope under various magnifications for histological detection of the general morphology of specimens. Representative images were acquired with a digital camera system using identical exposure settings across all specimens.

### Thiol and nucleotide content in blood

Blood (50 μL, WT n=6, PKU n=6) was immediately hemolyzed with 500 μL of ddH_2_O. Five hundred μL of this solution was then transferred to an Eppendorf microcentrifuge tube containing 750 μL of the precipitating solution (described above). The sample was kept on ice for 10 minutes, then centrifuged at 13,800 × g for 10 minutes at 4°C. The supernatant was frozen and stored at −20°C, while the pellet was discarded. Before HPLC injection, 15 μL of Na₂HPO₄ (0.3 M) and 7.5 μL of DTNB were added to 60 μL of the supernatant.

The mixture was used to quantify cysteine and GSH by HPLC. A BDS Hypersyl TM C18 column (150 mm x 4.6 mm, particle size 5 μm) (Thermo Scientific, USA) was used, coupled with a Teknokroma Tracer Excel ODS guard column (Teknokroma Analitica SA, Spain). The mobile phase consisted of 10 mM KH_2_PO_4_ solution, pH 6.0 (buffer A), and buffer A containing 60% (vol/vol) acetonitrile (buffer B). The elution conditions were as follows: 10 min at 100% buffer A, followed by 15 min an increase to 100% buffer B; this condition was maintained for 5 min. The gradient was returned to 100% buffer A over 3 min. The flow rate was 1 ml/min, the injection volume was 25 μl, and detection was at 330 nm using an HPLC Vanquish (Thermo Scientific, USA).

Blood (250 μL) was diluted with cold 0.5 M KOH (1:1 v/v) for basic extraction, vortexed, and incubated on ice for 5 minutes, then 250 μL of cold ddH_2_O was added. This suspension was centrifuged using an Amicon^®^ Ultra-4 (Merck Millipore, Ireland) at 4,000 × rpm for 20 minutes at 4 °C, and the procedure was repeated three times. The eluted solution was frozen and stored at −80 °C. Before HPLC injection, the pH was adjusted to 6.5 with 1 M KH_2_PO_4_. Nucleotide separation was performed as described in the previous HPLC section.

Thiols and nucleotide quantifications were normalized on hemoglobin concentration (mg/mL) determined with Drabkin’s reagent. Briefly, 10 μL hemolyzed blood was combined with 990 μL Drabkin’s solution [100 mg/L NaCN, 300 mg/L K_3_Fe (CN)_6_], and incubated for 10 min at RT in the dark. Absorbance was measured at 540 nm on the spectrophotometer.

### Statistical analyses

Statistical and graphical analyses were conducted using GraphPad Prism 11.0.0 (USA). Normality was assessed using the Shapiro-Wilk test, and differences between experimental groups were evaluated using Student’s *t*-test with Welch’s or Mann-Whitney *post hoc* correction unless otherwise stated. Results were considered statistically significant at *p* < 0.05. Quantitative image analyses were performed using ImageJ (USA).

## Results

Neurological aspects of PAH deficient PKU have been investigated for decades. Although neurotoxicity is the hallmark of PKU pathology and elevated Phe levels interfere with neurotransmitter synthesis, synaptic maturation, and overall metabolic homeostasis in the brain (Dobrowolski et al., 2016), the mechanism underlying this neurotoxicity remains unclear. In this work, we have investigated the underlying biochemical and molecular mechanisms by combining metabolomic and proteomic analyses and comparing the brains of WT and PKU mouse groups.

To obtain a global view of the molecular alterations associated with PAH deficiency, label-free proteomic and untargeted metabolomic datasets from WT and PKU brains were analyzed through a multi-omics workflow. This approach allowed us to identify individual proteins and metabolites altered by the disease and to define coordinated pathway-level changes supported by both molecular layers. Untargeted metabolomic analysis revealed a broader disease-associated metabolic shift. Overall, 340 annotated metabolic features met the predefined statistical and fold-change thresholds. Among these, 192 metabolites were increased, and 148 were decreased in PKU brains relative to WT. (Figure 1A), indicating extensive metabolic remodeling. As expected, the strongest metabolic alteration involved Phe-derived and phenylketone-related metabolites, consistent with defective PAH activity and accumulation of Phe catabolic derivatives.

**Figure 1.**
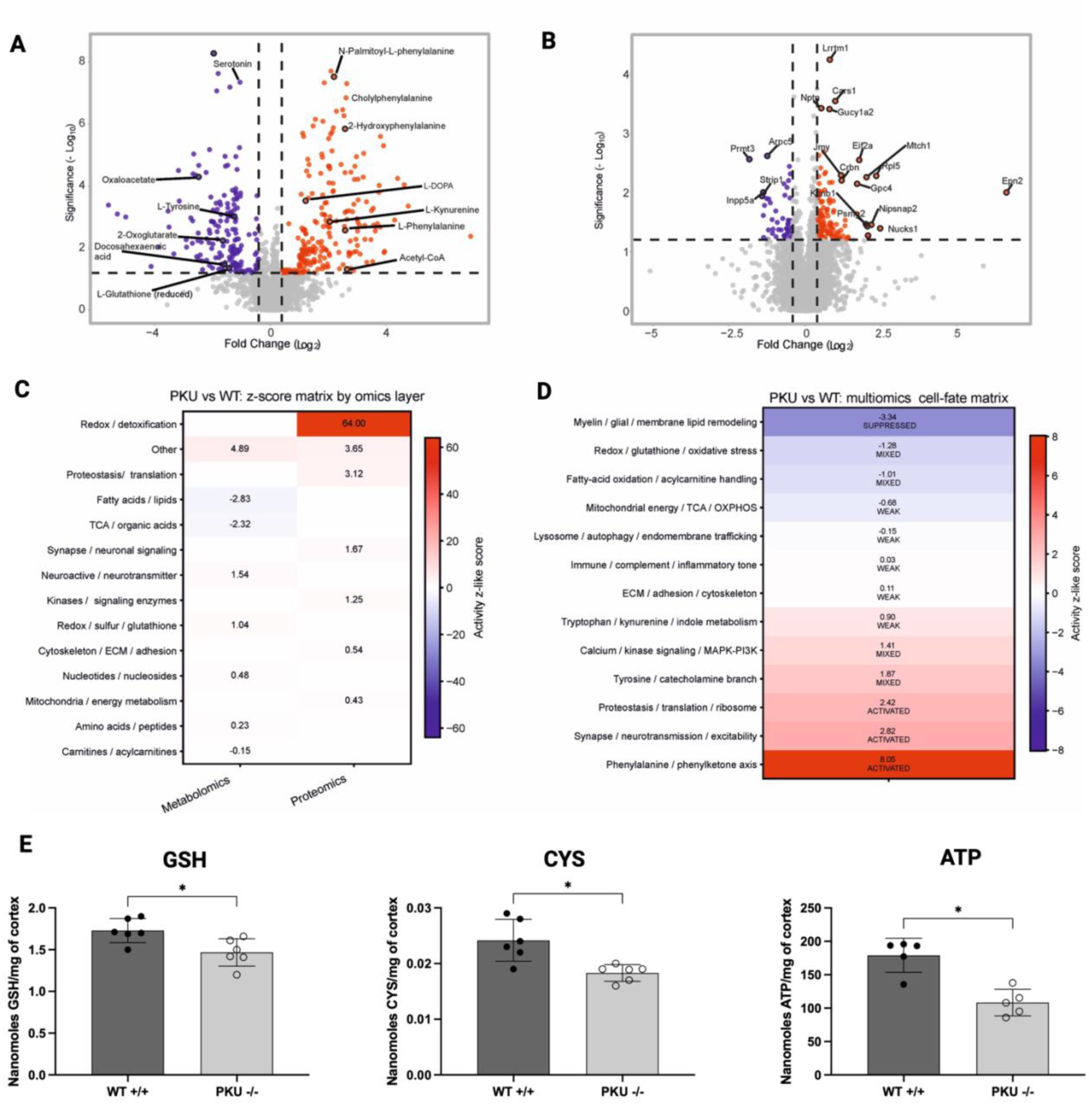
Proteomic, metabolomic, and integrated multi-omics remodeling in PAH^enu2^ brains. **(A)** Volcano plot of untargeted metabolomic changes in PAH^enu2^ brains relative to WT, showing increased Phe-related metabolites and decreased metabolites associated with tyrosine metabolism, redox balance, and central carbon metabolism. **(B)** Volcano plot of label-free proteomic changes in PKU brains relative to WT, highlighting significant alterations in proteins related to proteostasis, translation, neuronal signaling, and stress adaptive responses. **(C)** Z-score matrix showing pathway-level activity. Metabolomic and proteomic contributions are presented separately, highlighting specific alterations in redox and detoxification, proteostasis, the TCA cycle, and lipid-related pathways. **(D)** Integrated cell-fate matrix combining metabolomic and proteomic evidence into a single activity state. The Phe axis was the most strongly activated integrated module, whereas myelin glial remodeling was suppressed. Redox oxidative stress showed a mixed integrated pattern, consistent with pathway remodeling rather than uniform activation or suppression. **(E)** HPLC quantification of reduced glutathione (GSH), cysteine (CYS), and ATP levels in WT and PKU brain tissue. Data are expressed as mean ± SD normalized to mg of cortex (* < 0.05) (n=6/group).

Beyond the canonical Phe signature, the metabolomic dataset showed alterations in neuroactive metabolites, neurotransmitter-related pathways, tyrosine and catecholamine metabolism, redox-, sulfur-, and glutathione-associated metabolites, lipid species, and TCA organic acid intermediates. Neuroactive and neurotransmitter-related metabolites showed an overall positive trend, supporting the effect of PAH deficiency on neuronal signaling-related metabolic pools. In contrast, fatty acid and lipid metabolites showed a negative directional score, suggesting altered membrane lipid homeostasis. TCA and organic acid metabolites were also reduced, indicating impairment of central carbon and energy-related metabolism. Together, these metabolomic data supported the presence of amino acid toxicity, neurotransmitter imbalance, lipid remodeling, redox perturbation, and bioenergetic stress in PKU brains.

Label-free proteomic analysis identified 4,520 quantified proteins in WT and PKU brains. 148 proteins were significantly altered in PKU relative to WT, including 108 that were increased and 40 that were decreased (Figure 1B). Functional grouping of the altered proteins highlighted stress-adaptive and proteostasis-related processes, with prominent contributions from proteins involved in protein synthesis, ribosome function, protein quality control, redox homeostasis and detoxification, neuronal signaling, cytoskeletal organization, and endomembrane trafficking. These findings provided the molecular rationale for the subsequent biochemical investigation of translation initiation, ubiquitination, proteasome assembly and activity, autophagy-related markers, and ER-associated protein folding stress.

After separate analysis of the two molecular layers, metabolomic and proteomic results were integrated to identify coordinated pathway-level alterations. The integrated activity z-score analysis and the ON/OFF cell fate matrix (Figures 1C and 1D) revealed, apart from the Phe axis, activation of the synapse, neurotransmission, and excitability module, supported by altered neuroactive and amino acid-related metabolites, together with protein evidence of neuronal excitability and synaptic function. In parallel, the tyrosine and catecholamine branch showed a mixed or rewired profile rather than uniform activation or suppression, suggesting redistribution of neurotransmitter precursor and derivative pools. These findings indicate that PAH deficiency also affects broader neurochemical and neuronal signaling pathways.

Mitochondrial and energy metabolism showed a more complex pattern. The z-score matrix indicated a negative contribution of TCA metabolites, consistent with reduced central carbon intermediates. However, in the integrated cell-fate matrix, the mitochondrial energy/TCA/OXPHOS module showed only a weak negative score, suggesting partial pathway impairment rather than a uniformly suppressed mitochondrial program. This pattern is consistent with a model in which the metabolomic layer captures a deficit in energetic intermediates, whereas the proteomic layer may reflect compensatory or adaptive responses in mitochondrial and energy-associated proteins. Thus, the integrated data support the presence of bioenergetic stress in PKU brains, which was subsequently addressed by targeted biochemical analyses.

Redox metabolism was prominently affected but showed a mixed directional profile. The z-score matrix showed a strong proteomic contribution of redox pathways and a positive metabolomic contribution of redox, sulfur, and glutathione-related metabolites. Conversely, the integrated redox glutathione oxidative stress module in the cell-fate matrix was classified as mixed, indicating that redox homeostasis was remodeled rather than simply activated or suppressed. Accordingly, the integrated multi-omics signature provided the rationale for the subsequent evaluation of GSH, cysteine, antioxidant enzyme activity, and redox-sensitive pathways.

Lipid- and membrane-associated pathways were among the most consistently decreased components of the integrated analysis. The fatty acid and lipid metabolite family showed a negative activity score, and the myelin glial lipid remodeling module was the most strongly suppressed in the cell-fate matrix. Since membrane lipids are essential for myelin organization, synaptic vesicle dynamics, and neuronal membrane integrity, this suppressed lipid-associated signature supports the hypothesis that chronic hyperphenylalaninemia may affect structural and functional properties of neural membranes (Bregalda et al., 2023).

The integrated proteostasis and translation module was classified as activated in the cell fate matrix, in agreement with the proteomic evidence. This finding indicates that the PKU brain phenotype is associated with increased engagement of protein synthesis, turnover, and quality control pathways. Additional modules related to calcium, kinase signaling, lysosome and autophagy, endomembrane trafficking, and ECM showed weak or near-neutral integrated activity scores. Their inclusion in the integrated signature suggests broader cellular remodeling involving intracellular trafficking, structural organization, and signaling adaptation.

Overall, the combined proteomic, metabolomic, and integrated pathway analyses identified a coordinated molecular phenotype in PKU brains. These data suggest that brain pathology is not restricted to Phe accumulation but involves interconnected alterations in metabolism, oxidative stress, membrane function, and protein quality control. Based on this integrated molecular signature, subsequent analyses focused on validating redox imbalance, antioxidant enzyme activity, proteasome assembly and activity, autophagy markers, ER stress-related proteins, and systemic energetic redox parameters. These data may complement a previous metabolomic study by Dobrowolski and colleagues (Dobrowolski et al., 2022a), which suggested dysfunction of respiratory chain complex I in PKU brain.

The drop in GSH and cysteine content was further confirmed and quantified by HPLC, along with OXPHOS dysfunction, as monitored by ATP production (Figure 1E). Since metabolite alterations were identified as linked to a local oxidative stress environment, the activity of other redox enzymes, such as G6PD, SOD, and CAT, was also evaluated in PKU brains (Figure 2). G6PD, which converts glucose-6-phosphate to 6-phosphogluconolactone, showed a non-statistically significant trend toward increased activity, possibly reflecting a compensatory mechanism of the pentose phosphate pathway (PPP) aimed at sustaining NADPH production under conditions of impaired redox homeostasis (Yang et al., 2021). CAT and SOD provide the primary defense against ROS by catalyzing the dismutation of O^2•−^ to H_2_O_2_, which is then reduced to O_2_ and H_2_O (Bortoluzzi et al., 2021). First, SOD converts superoxide radicals into H_2_O_2_, which is then neutralized by CAT, producing H_2_O and O_2_. In our settings, both CAT and SOD enzyme activities were negatively affected by the disease, consistent with recent reports in a PKU rat hippocampus model (Bortoluzzi et al., 2019b; Çiçek et al., 2024).

**Figure 2.**
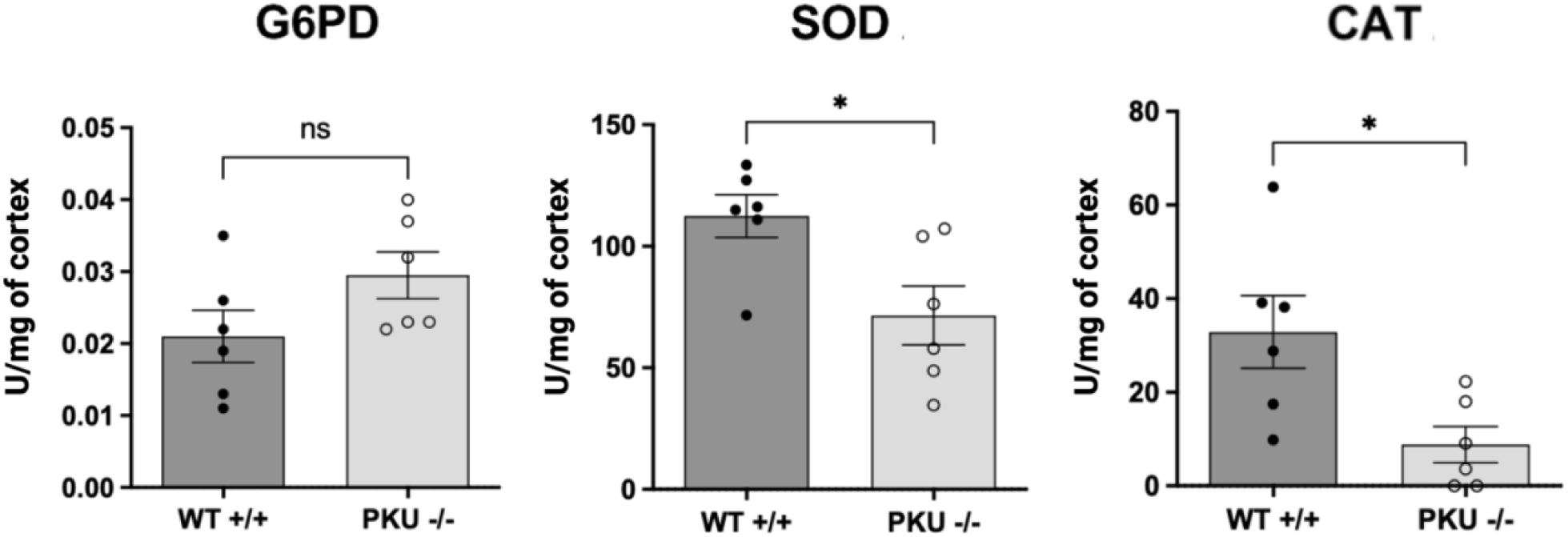
Antioxidant enzyme activities in WT and PAH^enu2^ mouse brains. Activities of glucose-6-phosphate dehydrogenase (G6PD), superoxide dismutase (SOD), and catalase (CAT) were measured in brain homogenates from WT and PKU mice. Data are expressed as mean ± SD normalized to mg of cortex (* < 0.05) (n=6/group).

To deepen the characterization of the redox status in the brains of PKU mice and to evaluate the potential activation of inflammatory pathways, gene expression in the *cerebrum* was performed, specifically for genes involved in i) oxidative response, i.e, *CHAC1* (γ-glutamyl cyclotransferase), *HMOX-1* (heme oxygenase-1), *GCLC*/*GCLM* (glutamate cysteine ligase C and M), *NQO1* (NAD[P]H dehydrogenase quinone-1), *NRF2* (Nuclear factor erythroid 2-related factor 2) and ii) inflammatory NF-kB and AP-1 downstream markers, i.e. IL-6 (interleukin-6), TNF-α (tumor necrosis factor-α), MCP-1 (monocyte chemoattractant protein-1) (Figure 3). Given that none of the above genes were modulated in PKU mice versus WT, a concomitant western blot analysis of the upstream transcription factors Nrf2 and the phosphorylated NF-kB subunit p65 confirmed the absence of activation of those factors (not shown), thereby supporting the absence of both anti-inflammatory gene response and activation of Nrf2-target genes.

**Figure 3.**
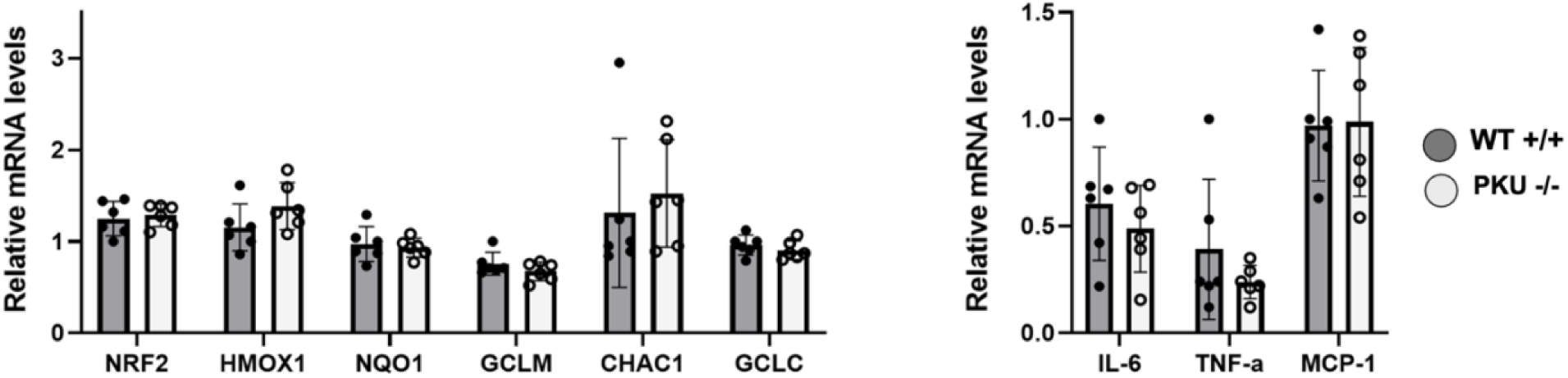
Gene expression of oxidative and inflammatory markers in WT and PKU brains. RT-PCR analyses of *CHAC1* (γ-glutamyl cyclotransferase), *HMOX-1* (heme oxygenase-1), *GCLC*/*GCLM* (glutamate cysteine ligase C and M), *NQO1* (NAD[P]H dehydrogenase quinone-1), *NRF2* (Nuclear factor erythroid 2-related factor 2, IL-6 (interleukin-6), TNF-α (tumor necrosis factor-α), MCP-1 (monocyte chemoattractant protein-1). Data are expressed as fold change relative to the control (mean ± SD). (n=6/group).

The possibility that protein translation was indeed increased in PKU mice, as suggested by -omics analyses, was further investigated by western immunoblotting of brain lysates. The results showed that phosphorylation of the eukaryotic initiation factor 2a ([p]-eIF2a) declined relative to its inactive counterpart (eIF2a), further supporting an enhancement of translation (Figure 4A). Since cellular proteins are constantly undergoing synthesis and degradation (Ross et al., 2020), we investigated the protein degradation machinery.

**Figure 4.**
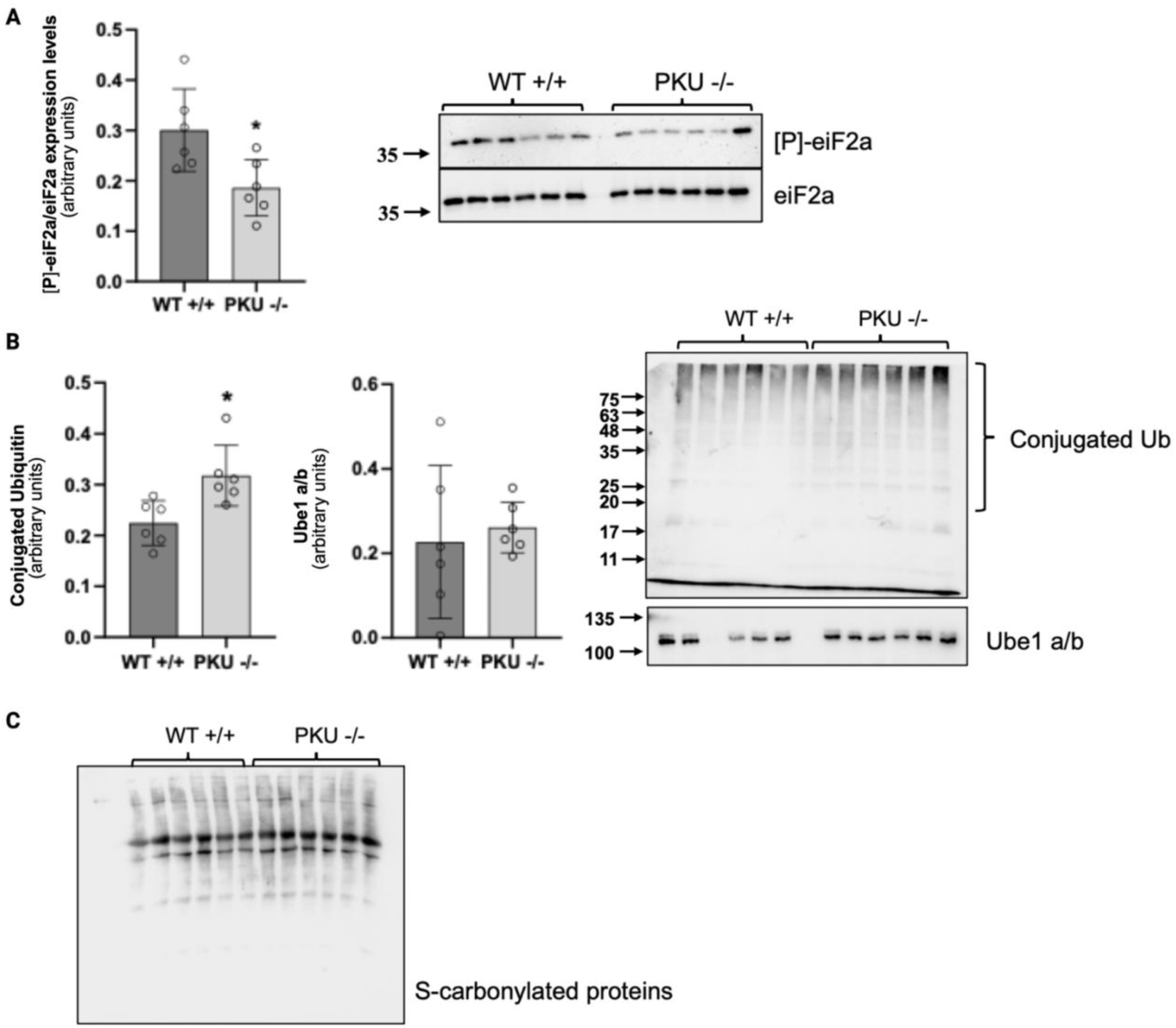
Protein translation and Ub-related protein degradation pathways in PAH^enu2^ brains. **(A)** Western blot analysis of phosphorylated eIF2α and total eIF2α expression in WT and PKU brain samples was performed by loading 20 µg of total protein lysate onto a 10% (w/v) acrylamide gel. The phosphorylation level of eIF2a was normalized to total eIF2a protein levels. **(B)** Ub-conjugated proteins were detected by Western blot analysis using an anti-Ub antibody. Five micrograms of total protein were separated by SDS–PAGE on a 14% (w/v) acrylamide gel. UBE1 expression was evaluated using an anti-UBE1a/b antibody, which detects endogenous levels of total UBE1a and UBE1b proteins. Twenty micrograms of total protein were separated by SDS–PAGE on a 7% (w/v) acrylamide gel. **(C)** Protein carbonylation in WT and PKU brains was assessed using an OxyBlot Protein Oxidation Detection Kit. Five micrograms of protein derivatized with DNPH were separated by SDS–PAGE on a 12% (w/v) acrylamide gel. Results are presented in the graphs as mean ± SD (*p < 0.05). Total protein staining was used as a loading control. (n=6/group).

Gene expression analysis of the stress-induced Ub genes *UBB* and *UBC* showed no modulation (not shown). Nevertheless, western immunoblotting of brain extracts revealed an accumulation of Ub-conjugated proteins in PKU mice. Since the level of protein ubiquitination reflects the rate of Ub conjugation and deconjugation, as well as protein degradation by the 26S proteasome, the levels of Ube1 and proteasome expression and activity were subsequently examined. Ube1, the first enzyme in the Ub-conjugation cascade, showed increased expression in PKU (Figure 4B). Despite previous data highlighting an imbalance in the redox status, no difference was observed between groups in carbonylated proteins, a marker of protein oxidation (Figure 4C).

By contrast, proteasome subunit expression did not change, except for increased levels of the 19S lid component Rpt1 in PKU mice compared to WT (Figure 5).

**Figure 5.**
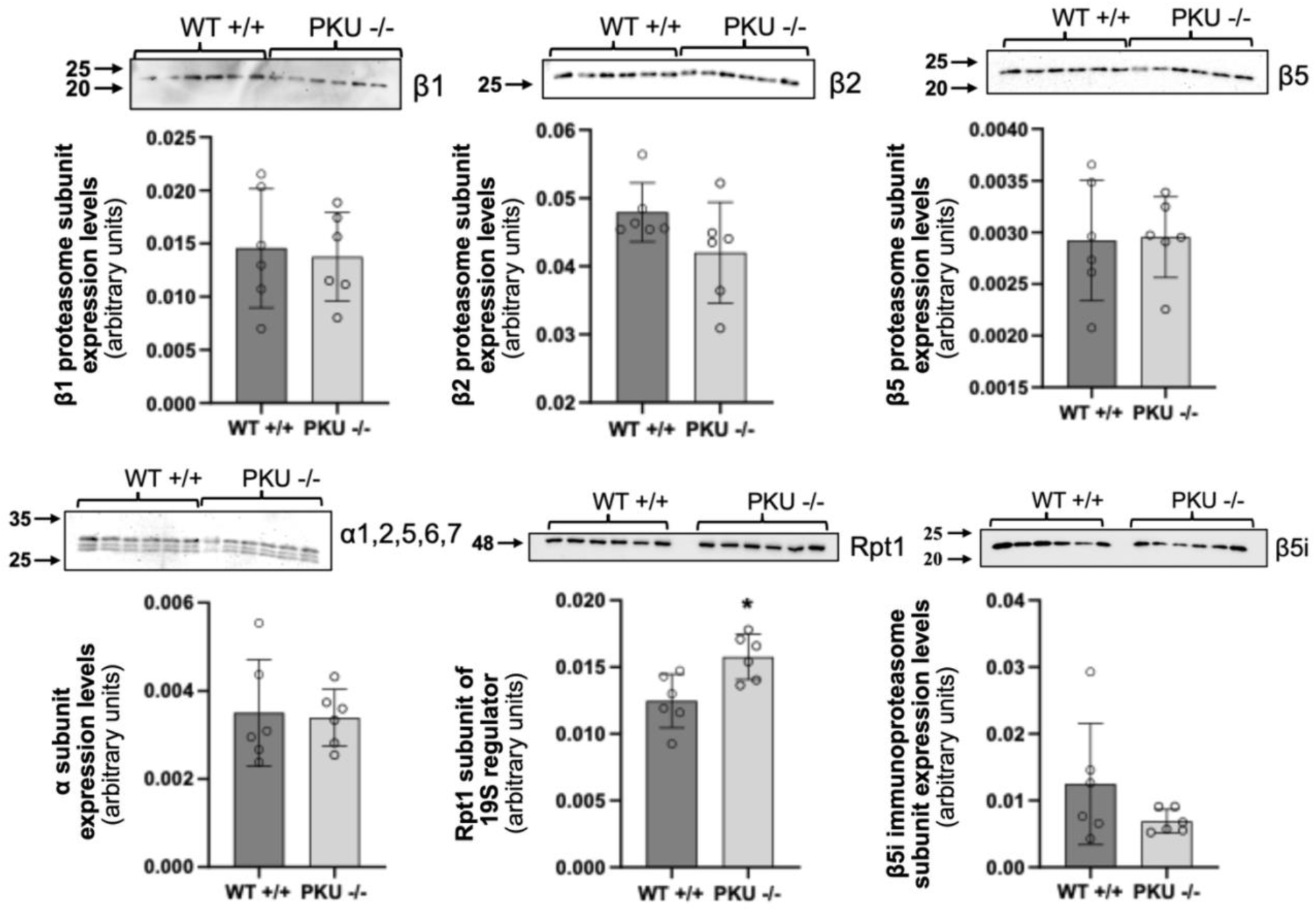
Expression of proteasome subunits in WT and PAH^enu2^ mouse brains. Representative Western blot analysis and relative quantification of proteasome subunits (panα, β1, β2, and β5), the immunoproteasome subunit β5i, and the Rpt1 subunit of the 19S regulatory particle. Two micrograms of total protein were separated by SDS–PAGE on a 12% (w/v) acrylamide gel for constitutive proteasome subunits, whereas 10 µg of total protein were loaded for detection of the β5i immunosubunit. Ten micrograms of lysates were separated on a 14% (w/v) acrylamide gel for the Rpt1 subunit of the 19S particle. Total protein staining was used as a loading control. Densitometric values of the immunoreactive bands were normalized to total protein content and are reported in the graphs. Data are presented as mean ± SD (*p < 0.05) (n=6/group).

Interestingly, chymotrypsin-like, trypsin-like, and caspase-like activities were significantly elevated in PKU mice, as measured in a cell-free system using fluorogenic substrates (Figure 6A). Consistent with this observation, PKU brains contained higher levels of active 19S-capped complexes, as revealed by native PAGE analyses followed by in-gel activity and western immunoblotting (Figure 6B).

**Figure 6.**
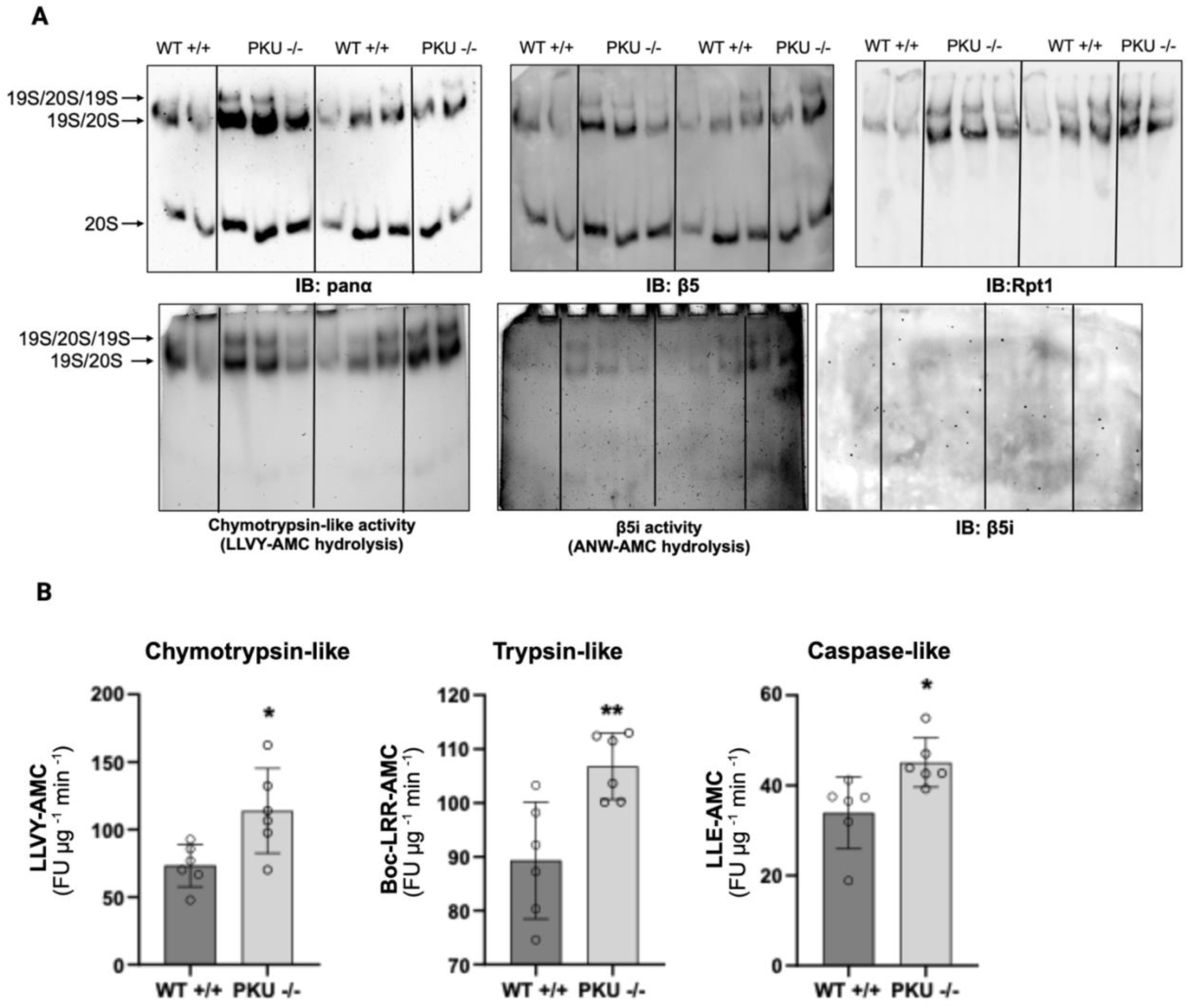
Proteasome activity and assembly. **(A)** Representative immunoblot (IB) images of native PAGE analysis of proteasome complexes. Twenty micrograms of native protein extracts were separated on a 4% (w/v) acrylamide gel under native conditions in the presence of ATP and subjected to Western blot analysis using specific antibodies against α subunits, β5, β5i, and Rpt1. Chymotrypsin-like and β5i-associated activities were visualized by in-gel peptidase activity assay using the fluorogenic substrates sLLVY-AMC and ANW-AMC, respectively. Following separation of proteasome complexes by native PAGE, gels were incubated with the substrates, and fluorescent signals were detected using a ChemiDoc Imaging System under UV light. Immunoproteasome activity was minimally detectable. **(B)** Proteasome activities were measured in cell-free extracts prepared under native conditions using the fluorogenic peptide substrates sLLVY-AMC, Boc-LRR-AMC, and LLE-AMC for chymotrypsin-like, trypsin-like, and caspase-like activities, respectively. Activity values are expressed as UF/min/µg protein and are reported in the graphs (FU, fluorescence units). Data are presented as mean ± SD (\*\**p* < 0.01; \**p* < 0.05) (n=6/group).

The immunoproteasome (detected by hydrolysis of the fluorogenic substrate ANW-AMC and b5i staining) was barely detectable, consistent with the absence of pro-inflammatory stimuli in the brain tissue. Overall, these data indicate increased protein turnover in this disease model, which may reflect a higher load of oxidatively damaged or misfolded proteins that require degradation via the UPS. In parallel to the UPS, autophagy was explored as the other primary mechanism for protein degradation. To this end, the levels of proteins degraded during the process and those maintained during autophagy were analyzed. This approach allows assessment of autophagic flux in tissues where lysosomal activity cannot be blocked (Humbert et al., 2020). Autophagosome markers LC3-II, which is degraded within mature autophagosomes and autolysosomes [33], and its precursor LC3-I were both expressed at lower levels in PKU compared to WT. Moreover, the LC3-/d LC3-I ratio was also decreased. Similarly, levels of another autophagy substrate, p62/SQSTM1, showed a decreasing trend, although it was not statistically significant (Figure 7). By contrast, the intracellular content of the non-substrate protein of autophagy, ATG5, was unchanged (Tutas et al., 2025) (Figure 7). Accordingly, specific heat shock protein (HSP), thPA8, known to regulate autophagy (Dong et al., 2022), was found to be overexpressed in PKU brain (Figure 7). Altogether, this evidence suggests higher autophagic activity in PKU brains.

**Figure 7.**
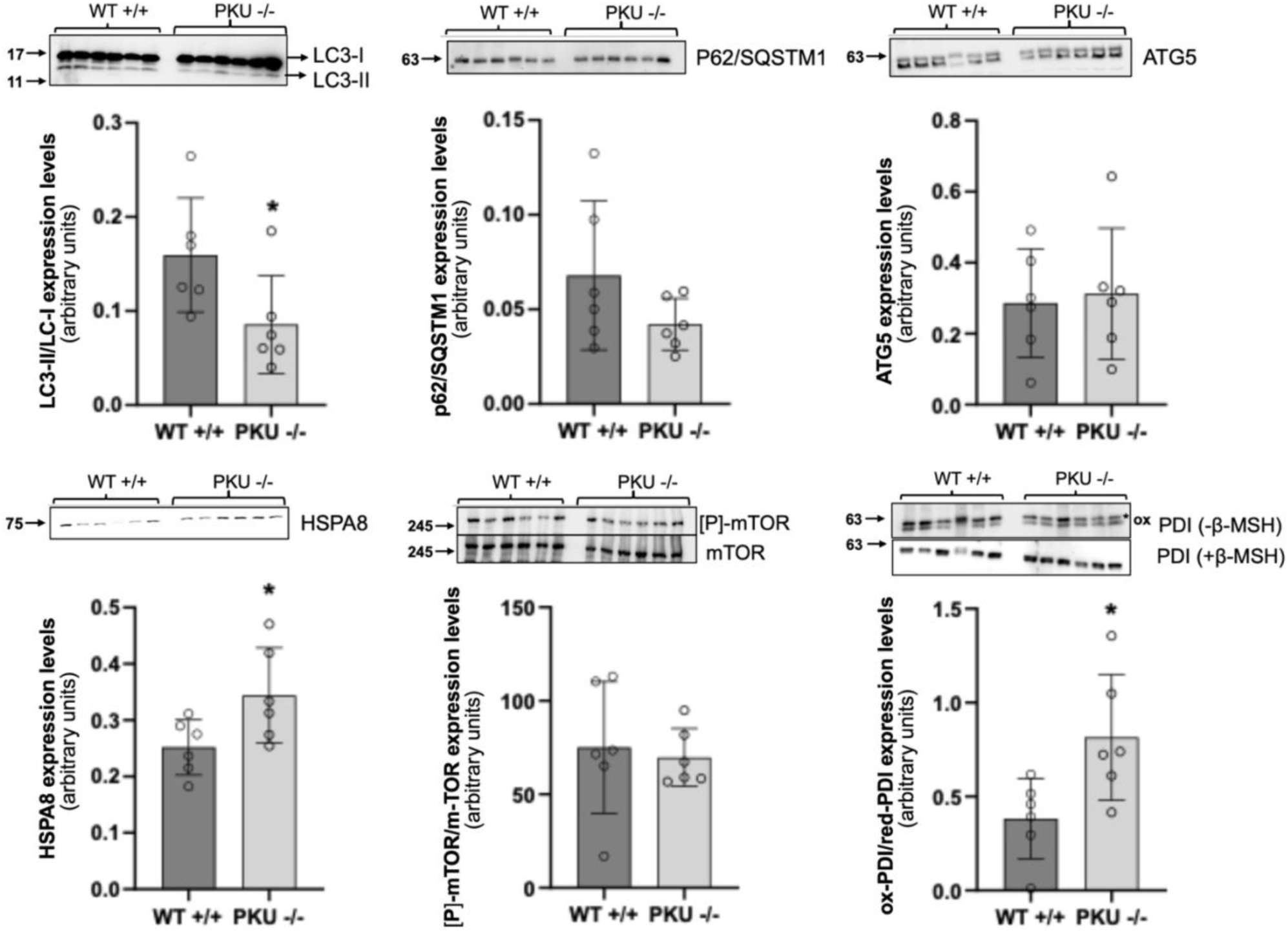
Autophagy markers and ER stress-related proteins. Western blot analysis of autophagy-associated proteins LC3-I/LC3-II, p62/SQSTM1, ATG5, HSPA8, and [P]-mTOR/mTOR was performed in WT and PKU brain lysates. LC3-I/LC3-II were analyzed by loading 10 µg of total lysate onto a 14% (w/v) acrylamide gel, and the results are shown in the graph as the LC3-II/LC3-I ratio. ATG5, HSPA8, and p62/SQSTM1 were evaluated by loading 20 µg of total lysates onto 10% (w/v) acrylamide gels. Phosphorylated mTOR ([P]-mTOR) and total mTOR were analyzed by loading 25 µg of total lysates onto a 6% (w/v) acrylamide gel. PDI oxidation status was assessed under nonreducing conditions, in the absence of β-mercaptoethanol (β-MSH), by loading 50 µg of total lysates onto an 8% (w/v) acrylamide gel, whereas total PDI levels were evaluated under reducing conditions by loading 20 µg of total lysates onto an 8% (w/v) acrylamide gel. Total protein staining was used as a loading control for LC3-I/LC3-II, HSPA8, ATG5, and nonreduced PDI, whereas β-actin was used as a normalization control for p62/SQSTM1 and reduced PDI. The phosphorylation level of [P]-mTOR was normalized to total mTOR expression. Quantification of normalized values for each target is reported in the respective graphs as the mean ± SD (n = 6 per group). Representative immunoblots are shown (\**p* < 0.05).

The mTOR signaling pathway is one of the main pathways regulating autophagy [37]. However, in PKU, the ratio of phospho-mTOR (p-mTOR) to mTOR did not change, indicating that mTOR was not activated (Figure 7), suggesting an alternative regulatory mechanism.

To determine the consequences of increased protein turnover, we next evaluated UPR activation and HSP90 expression, since they reflect the cellular response to an increased burden of misfolded proteins. In our settings, neither Hsp90 nor ER stress sensors such as IRE1 and ATF6 were modulated. Accordingly, BIP (Grp78) expression levels did not change in PKU brains (Figure S1). Sustained maintenance of specific redox conditions for the formation of correct disulfide bonds inside the ER requires equilibrium between the oxidized and reduced states of protein disulfide isomerases (PDIs) (Andreu et al., 2012). We detected higher levels of oxidized PDI in PKU than in WT, suggesting a perturbation of the local oxidative environment that did not reach the threshold for UPR activation. (Figure 7).

To assess whether the molecularly identified biochemical alterations were accompanied by morphological changes, a qualitative histological examination of PKU mouse brains was performed. Histological evaluation revealed structural alterations broadly consistent with the molecular signatures identified in the proteostasis and redox analyses. H&E staining showed neuronal rarefaction, shrinkage of neuronal cell bodies, and occasional pyknotic nuclei in both the cortex and cerebellum (Supplementary Figures S2 and S3). In addition, focal inflammatory areas with perivascular and parenchymal lymphocyte infiltration were observed (Supplementary Figure S4). These findings may be consistent with localized inflammatory changes. Overall, these observations provide morphological evidence that can be considered alongside the molecular and metabolomic data presented in this study.

Although preliminary, these observations support the notion that metabolic and proteostatic alterations may eventually translate into compensatory changes within vulnerable brain regions.

To further assess whether a systemic deficiency in antioxidant defenses was present, we measured circulating antioxidant markers. Data from PKU patients obtained from peripheral tissues, especially blood, showed decreased antioxidant response, possibly due to dietary restriction of micro- or macronutrients with antioxidant properties (Rocha and Martins, 2012). In this work, redox and bioenergetic parameters were measured in whole-mouse blood to determine whether redox dysregulation contributes to the pathophysiology of tissue damage in the absence of Phe restriction. The HPLC results showed that both ATP and ADP decreased (Figure 8). Erythrocytes do not directly synthesize ADP from adenosine but instead obtain it from ATP or AMP; therefore, Phe could inhibit pyruvate kinase, likely reducing ATP and, consequently, lowering ADP levels. On the other hand, NADPH was increased, although this couldn’t be correlated to its production through the pentose phosphate pathway because G6PD activity didn’t vary. NADPH content correlated with reduced erythrocyte GPX and GR activities, as reported by Sirtori *et al*. (Sirtori et al., 2005), which, in turn, could be responsible for the decreased GSH levels. Being GSH the main antioxidant arsenal of a cell vulnerable to oxidative damage and, being the *savage* pathway from GSSG to GSH limited by GR activity, *de novo* synthesis using the rate-limiting amino acid cysteine, depleted RBC’s cysteine reservoir. These data suggest that the observed cerebral oxidative damage could reflect a whole-body redox imbalance rather than an isolated, tissue-specific phenomenon.

**Figure 8.**
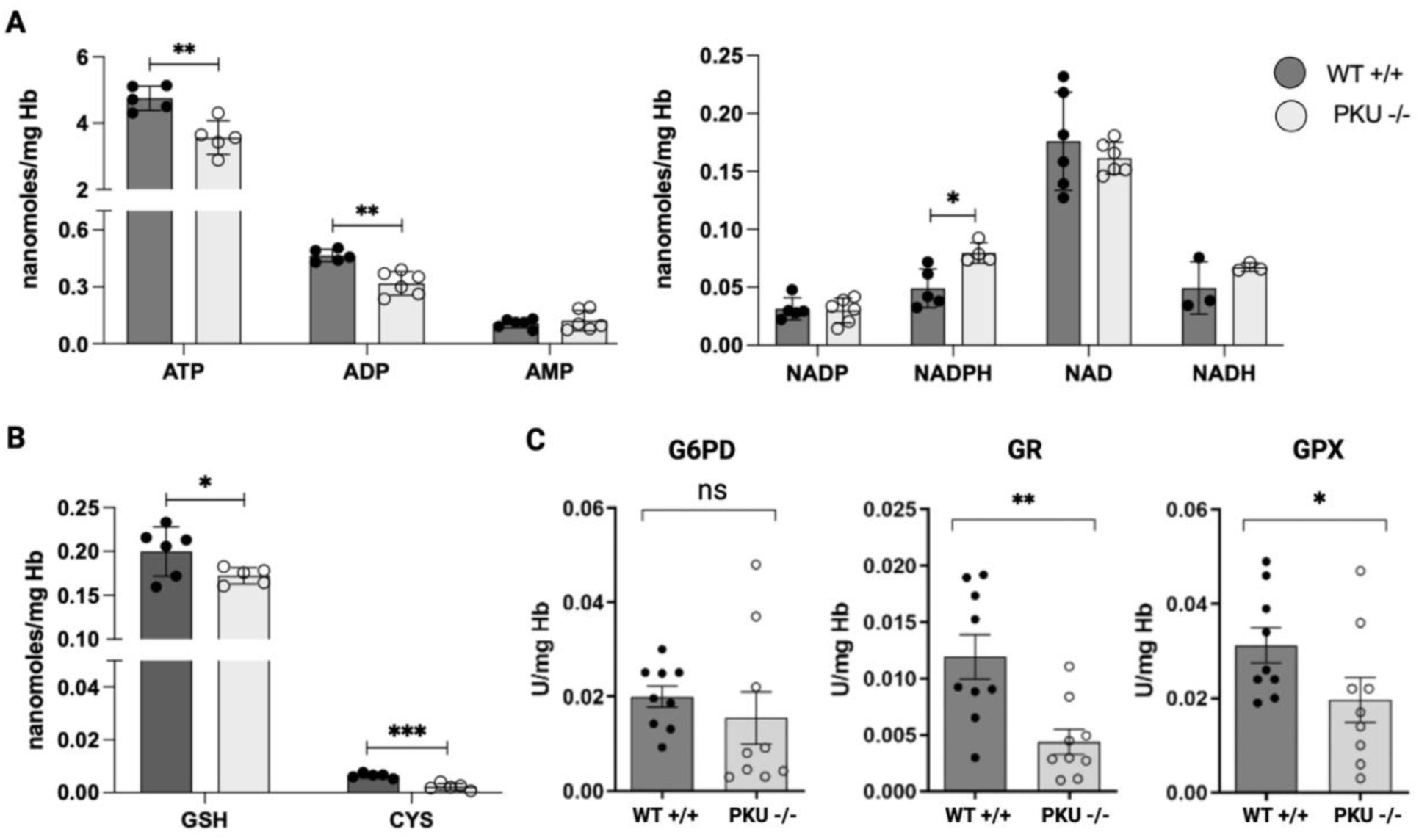
Systemic redox and energetic alterations in whole blood from PAH^enu2^ mice. HPLC quantification of **(A)** ATP, ADP, AMP, NAD, and NADH levels from WT and PKU mice; **(B)** reduced glutathione (GSH) and cysteine contents in whole blood (n=6/group). **(C)** Activities of glutathione peroxidase (GPX), glutathione reductase (GR), and glucose-6-phosphate dehydrogenase (G6PD) measured in blood hemolysates (n=9/group). Data are expressed as mean ± SD normalized on hemoglobin (Hb) values (* < 0.05; ** < 0.01).

## Discussion

Processes such as alterations in protein synthesis, oxidative stress, and changes in bioenergetics may play crucial roles in the development of neurological symptoms in PKU. Using an in-house mouse model of PKU, the BT-BR PAH^enu2^, which carries a loss-of-function mutation in the murine PAH enzyme, we investigated whether there are differences in the molecular mechanisms of homeostasis maintenance between brains derived from WT and PKU mice in terms of redox and proteostatic balance, given that high levels of Phe have been shown to cause indirect neurodegeneration due to loss of synaptic plasticity and metabolic, and structural alterations (Schlegel et al., 2016). Metabolomic profiling revealed an oxidative environment in PKU brains, with decreased GSH and elevated GSSG levels, along with alterations in GSH transferases and peroxiredoxins.

The metabolomic profile additionally revealed substantial alterations in neurotransmission-related pathways, including dysregulation of amino acid metabolism and GABA-associated signaling. These findings were consistent with previous evidence demonstrating impaired monoaminergic neurotransmission and synaptic maturation in PKU [2,41]. Furthermore, the observed lipid remodeling may alter membrane fluidity and synaptic vesicle dynamics, thereby exacerbating neuronal dysfunction. The histologically observed neuronal rarefaction and leukocyte infiltration may reflect oxidative imbalance and enhanced protein degradation leading to disruption of neuronal homeostasis, and ultimately structural damage.

Nevertheless, PKU mice did not exhibit activation of either the antioxidant response (i.e., Nrf2) or pro-inflammatory mechanisms (i.e., NF-κB). Absent modulation of Nrf-2 activity is corroborated by unchanged expression of its target genes GCLC, GCLM, and the associated protein products, which in turn suggest that GSH decrease is not due to impaired synthesis. Still, it could be correlated with altered function/expression of antioxidant enzymes such as glutathione peroxidase and reductase, as determined in blood. Dysregulation of Nrf2 signaling has been linked to several diseases associated with oxidative stress and inflammation, including neurological disorders (Cuadrado, 2022). However, we could postulate that these mice could activate other antioxidant defense mechanisms to avoid oxidative damage and neuroinflammation, such as glutaredoxins and thioredoxins, and have already been demonstrated to mediate neuroprotection against oxidative stress in neurodegenerative diseases (Seco-Cervera et al., 2020).

Alteration of redox homeostasis is confirmed by NADPH accumulation and decreased cysteine-GSH levels in the blood of mice. GSH becomes an essential antioxidant for cells that are more vulnerable to oxidative damage, such as RBCs, even though RBCs inherently contain high levels of GSH. Reduced GSH levels in cells could be counteracted by *de novo GSH* synthesis, as revealed by significant cysteine consumption. Concomitantly, a possible impairment of GSH peroxidase activity or expression would lead to an increase in NADPH levels.

Moreover, we found that the proteolytic system was more active in the brains of PKU mice than in those of WT counterparts. This increase involved both the Ub-proteasome system and autophagy/lysosomal systems (UPS and ALS). To date, no studies have examined PKU-associated changes in the functional activity and composition of the proteasome pool in the mouse cerebral cortex. Hereby, the expression of proteasome genes, their subunit composition, and the activity of individual forms (including those containing immune subunits) were thoroughly investigated. Interestingly, increased proteasome activity in PKU brains was not due to increased expression of proteasome components (except for Rpt1 of the 19S RP), but rather to higher assembled proteasome complexes, suggesting a boost in the proteasomal machinery. It has been suggested that proteasomal degradation of proteins yields free amino acids, which are then recycled for the synthesis of new proteins, or that the peptides are redirected into metabolic pathways to maintain cellular energy (Brennan et al., 2025). Moreover, in our model, the molecular chaperone HSPA8, which acts as a folding catalyst, ensuring the refolding of misfolded conformers, and as a controller that targets proteins for subsequent degradation (Bonam et al., 2019), was found to be highly expressed in PKU brains. HSPA8 is crucial for chaperone-mediated autophagy, supporting the push of the degradation machinery in our disease model. Taken together, these data suggest that chaperones and degradative pathways in the brains of PKU mice work synergistically to maintain proteostasis without triggering the ISR (integrated stress response), as shown by reduced phosphorylation of eIF2a, and to overcome endoplasmic reticulum folding quality control, as no markers of the UPR were found. However, PDI, a thiol-disulfide switch crucial for maintaining ER homeostasis and preventing the accumulation of misfolded proteins (Owegie et al., 2025), is predominantly in its oxidized form and could trigger UPR activation at a later stage. Misfolded or aberrant proteins require reduction of their disulfide bonds prior to cytosol translocation for degradation by the proteasome. PDI, in its reduced form, is involved in breaking disulfide bonds that otherwise stabilize misfolded proteins and impede their clearance (Owegie et al., 2025); therefore, the oxidized form present in our settings could mirror an enhanced degradation activity.

Proteasomal degradation is an ATP-dependent process tightly linked to cellular energy homeostasis, and neurons are particularly vulnerable to disturbances in protein quality control due to their high metabolic demand and limited regenerative capacity. In our model, enhanced degradation capacity occurred concomitantly with reduced ATP levels and altered Krebs cycle intermediates, suggesting that the proteostasis network operates under energetically unfavorable conditions. Such an increase in ATP-consuming proteolytic activity may reflect the necessity to preserve neuronal integrity by rapidly eliminating damaged or dysfunctional proteins generated under chronic hyperphenylalaninemia.

While components, such as the UPR in the ER and mitochondria, have been extensively characterized in several disease models (Kuzu et al., 2025), their interconnections remain unclear. Interactions between ER stress signaling and mitochondrial pathways, as well as cross-regulation between the ER, UPR, and stress responses, deserve further investigation. A more integrated understanding of these networks will lay the groundwork for developing targeted therapeutic strategies. Nevertheless, further research is needed to determine the potential impact of the Phe-dependent proteostasis network on brain structure and function. Although we demonstrated activation of proteolytic systems, functional studies assessing neuronal activity and behavioral outcomes were beyond the scope of this work. Future studies should investigate whether modulation of proteasome activity, autophagy, or redox metabolism may ameliorate neurological manifestations in PKU.

In conclusion, this work provides the first comprehensive characterization of proteasome assembly and activity in the PKU mouse brain and identifies proteostasis remodeling as a component of PKU neuropathology. Together with oxidative and metabolic alterations, enhanced activation of proteolytic pathways emerges as a central adaptive mechanism in response to chronic Phe-induced stress. These findings broaden the current understanding of PKU pathophysiology and may offer new perspectives for therapeutic strategies in neurometabolic disorders.

## Abbreviations

AP-1: Activator Protein-1
ATF4: Activating transcription factor 4
ATF6: Activating transcription factor 6
CAT: Catalase
CHAC1: γ-Glutamyl cyclotransferase
eIF2α: Eukaryotic translation initiation factor 2 α
ER: Endoplasmic reticulum
G6PD: Glucose-6-phosphate dehydrogenase
GCLC/GCLM: Glutamate cysteine ligase C and M
GSH: Reduced glutathione
GSH-Px: Glutathione peroxidase
GSSG: Oxidized glutathione
HMOX-1: Heme oxygenase-1
HSP90: Heat shock protein 90
IL-6: Interleukin 6
IRE1α: Inositol-requiring kinase 1α
MCP-1: Monocyte chemoattractant protein-1
NF-κB: Nuclear Factor kappa-light-chain-enhancer of activated B cells
Nrf2: Nuclear factor erythroid 2-related factor 2
NQO1: NAD[P]H dehydrogenase quinone-1
OxS: Oxidative Stress
PAH: Phenylalanine hydroxylase
PDI: protein disulfide isomerase
PERK: Protein kinase-like ER kinase
PHE: Phenylalanine
PPP: pentose phosphate pathway
PKU: Phenylketonuria
ROS: Reactive oxygen species
SOD: Superoxide dismutase
TNF-α: Tumor necrosis factor-α
TYR: Tyrosine
UB: Ubiquitin
UPS: Ubiquitin-proteasome system
UPR: Unfolded protein response
WT: Wild-type
XBP1: X-box binding protein 1

## Declaration of generative AI and AI-assisted technologies in the writing process

During the preparation of this work, the authors used an OpenAI chatbot in order to improve language and readability. The authors reviewed and edited the content as needed and took full responsibility for the content of the publication

## Acknowledgments

The authors gratefully acknowledge Dr. Sofia Masini for insightful support with the mouse study and helpful suggestions throughout. We also thank Claudia Scopa for the help in animal handling. Figures and graphical illustrations were created, in part, using BioRender.com.

We would also like to thank POR MARCHE FESR 2014/2020. Asse 1, OS 2, Azione 2.1 – Intervento 2.1.1 – Sostegno allo sviluppo di una piattaforma di ricerca collaborativa negli ambiti della specializzazione intelligente. Thematic Area: “Medicina personalizzata, farmaci e nuovi approcci terapeutici”. Project acronym: Marche BioBank www.marchebiobank.it. «The content of the paper is the sole responsibility of the authors and can under no circumstances be regarded as reflecting the position of the European Union and/or Marche Region authorities».

## CrediT authorship contribution statement

**Francesca Monittola:** Investigation, Data curation, Visualization, Review & editing. **Elena Perla:** Investigation, Data curation, Review & editing. **Debora Libetti:** Investigation, Data curation, Review & editing. **Antonella Antonelli**: Investigation, Data curation, Visualization, Review & editing. **Laura Graciotti:** Investigation, Data curation, Visualization, Review & editing. **Davide Torre:** Investigation, Data curation. **Francesca Pierigé:** Investigation, Resources. **Anastasia Ricci:** Investigation. **Mauro Magnani:** Review & editing. **Marzia Bianchi:** Data curation, Visualization, Review & editing. **Sara Biagiotti:** Data curation, Visualization, Review & editing. **Luigia Rossi:** Conceptualization, Formal analysis, Resources, Funding. **Michele Menotta:** Data curation, Visualization, Review & editing. **Alessandra Fraternale:** Conceptualization, Formal analysis, Resources, Data curation, Visualization, Review & editing. **Rita Crinelli:** Conceptualization, Methodology, Formal analysis, Resources, Data curation, Visualization, Review & editing. **Michela Bruschi:** Conceptualization, Formal analysis, Investigation, Data curation, Software, Writing original draft, review & editing, Visualization.

## Ethics approval and consent to participate

Not applicable

## Availability of data and materials

All data generated or analyzed during this study are included in this published article and the supplementary information file. The raw data and materials in the study are available from the corresponding authors on reasonable request.

## References

Andreu, C.I., Woehlbier, U., Torres, M., Hetz, C., 2012. Protein disulfide isomerases in neurodegeneration: from disease mechanisms to biomedical applications. FEBS Lett 586, 2826–2834. 10.1016/j.febslet.2012.07.023

Beutler E., 1984. Glutathione peroxidase, in: Red Cell Metabolism. A Manual of Biochemical Method. Grune & Stratton, Inc., pp. 65–66.

Bickel, H., Gerrard, J., Hickmans, E.M., 1953. Influence of phenylalanine intake on phenylketonuria. Lancet 265, 812–813. 10.1016/s0140-6736(53)90473-5

Bobak, Y., Kurlishchuk, Y., Vynnytska-Myronovska, B., Grydzuk, O., Shuvayeva, G., Redowicz, M.J., Kunz-Schughart, L.A., Stasyk, O., 2016. Arginine deprivation induces endoplasmic reticulum stress in human solid cancer cells. Int J Biochem Cell Biol 70, 29–38. 10.1016/j.biocel.2015.10.027

Bonam, S.R., Ruff, M., Muller, S., 2019. HSPA8/HSC70 in Immune Disorders: A Molecular Rheostat that Adjusts Chaperone-Mediated Autophagy Substrates. Cells 8, 849. 10.3390/cells8080849

Bortoluzzi, V.T., Brust, L., Preissler, T., de Franceschi, I.D., Wannmacher, C.M.D., 2019a. Creatine plus pyruvate supplementation prevents oxidative stress and phosphotransfer network disturbances in the brain of rats subjected to chemically-induced phenylketonuria. Metab Brain Dis 34, 1649–1660. 10.1007/s11011-019-00472-7

Bortoluzzi, V.T., Brust, L., Preissler, T., de Franceschi, I.D., Wannmacher, C.M.D., 2019b. Creatine plus pyruvate supplementation prevents oxidative stress and phosphotransfer network disturbances in the brain of rats subjected to chemically-induced phenylketonuria. Metab Brain Dis 34, 1649–1660. 10.1007/s11011-019-00472-7

Bortoluzzi, V.T., Dutra Filho, C.S., Wannmacher, C.M.D., 2021. Oxidative stress in phenylketonuria-evidence from human studies and animal models, and possible implications for redox signaling. Metab Brain Dis 36, 523–543. 10.1007/s11011-021-00676-w

Bregalda, A., Carducci, C., Viscomi, M.T., Pierigè, F., Biagiotti, S., Menotta, M., Biancucci, F., Pascucci, T., Leuzzi, V., Magnani, M., Rossi, L., 2023. Myelin basic protein recovery during PKU mice lifespan and the potential role of microRNAs on its regulation. Neurobiol Dis 180, 106093. 10.1016/j.nbd.2023.106093

Brennan, P.N., MacMillan, M., Manship, T., Moroni, F., Glover, A., Troland, D., MacPherson, I., Graham, C., Aird, R., Semple, S.I.K., Morris, D.M., Fraser, A.R., Pass, C., McGowan, N.W.A., Turner, M.L., Manson, L., Lachlan, N.J., Dillon, J.F., Kilpatrick, A.M., Campbell, J.D.M., Fallowfield, J.A., Forbes, S.J., 2025. Autologous macrophage therapy for liver cirrhosis: a phase 2 open-label randomized controlled trial. Nat Med 31, 979–987. 10.1038/s41591-024-03406-8

Bruschi, M., Masini, S., Biancucci, F., Piersanti, G., Canonico, B., Menotta, M., Magnani, M., Fraternale, A., 2025. Redox modulation via a synthetic thiol compound reshapes energy metabolism in endothelial cells and ameliorates angiogenic expression in a co-culture study with activated macrophages. Biochim Biophys Acta Gen Subj 1869, 130803. 10.1016/j.bbagen.2025.130803

Buttari, B., Tramutola, A., Rojo, A.I., Chondrogianni, N., Saha, S., Berry, A., Giona, L., Miranda, J.P., Profumo, E., Davinelli, S., Daiber, A., Cuadrado, A., Di Domenico, F., 2025. Proteostasis Decline and Redox Imbalance in Age-Related Diseases: The Therapeutic Potential of NRF2. Biomolecules 15, 113. 10.3390/biom15010113

Çiçek, Ç., Gök, M., Bodur, E., 2024. Rat PKU Model Display Gender-Based Neuroinflammatory Changes: Proinflamatuary Cytokines and Lipid Peroxidation. MMJ 11, 30–37. 10.47572/muskutd.1388547

Colomé, C., Artuch, R., Sierra, C., Brandi, N., Lambruschini, N., Campistol, J., Vilaseca, M.-A., 2003. Plasma thiols and their determinants in phenylketonuria. Eur J Clin Nutr 57, 964–968. 10.1038/sj.ejcn.1601631

Costa-Mattioli, M., Walter, P., 2020. The integrated stress response: From mechanism to disease. Science 368, eaat5314. 10.1126/science.aat5314

Cuadrado, A., 2022. Brain-Protective Mechanisms of Transcription Factor NRF2: Toward a Common Strategy for Neurodegenerative Diseases. Annu Rev Pharmacol Toxicol 62, 255–277. 10.1146/annurev-pharmtox-052220-103416

Dobrowolski, S.F., Lyons-Weiler, J., Spridik, K., Vockley, J., Skvorak, K., Biery, A., 2016. DNA methylation in the pathophysiology of hyperphenylalaninemia in the PAHenu2 mouse model of phenylketonuria. Mol Genet Metab 119, 1–7. 10.1016/j.ymgme.2016.01.001

Dobrowolski, S.F., Phua, Y.L., Sudano, C., Spridik, K., Zinn, P.O., Wang, Y., Bharathi, S., Vockley, J., Goetzman, E., 2022a. Comparative metabolomics in the Pahenu2 classical PKU mouse identifies cerebral energy pathway disruption and oxidative stress. Mol Genet Metab 136, 38–45. 10.1016/j.ymgme.2022.03.004

Dobrowolski, S.F., Phua, Y.L., Vockley, J., Goetzman, E., Blair, H.C., 2022b. Phenylketonuria oxidative stress and energy dysregulation: Emerging pathophysiological elements provide interventional opportunity. Mol Genet Metab 136, 111–117. 10.1016/j.ymgme.2022.03.012

Dong, Y., Li, T., Ma, Z., Zhou, C., Wang, X., Li, J., 2022. HSPA1A, HSPA2, and HSPA8 Are Potential Molecular Biomarkers for Prognosis among HSP70 Family in Alzheimer’s Disease. Dis Markers 2022, 9480398. 10.1155/2022/9480398

Eichinger, A., Danecka, M.K., Möglich, T., Borsch, J., Woidy, M., Büttner, L., Muntau, A.C., Gersting, S.W., 2018. Secondary BH4 deficiency links protein homeostasis to regulation of phenylalanine metabolism. Hum Mol Genet 27, 1732–1742. 10.1093/hmg/ddy079

Embury, J.E., Reep, R.R., Laipis, P.J., 2005. Pathologic and immunohistochemical findings in hypothalamic and mesencephalic regions in the pah(enu2) mouse model for phenylketonuria. Pediatr Res 58, 283–287. 10.1203/01.PDR.0000170000.78670.44

Ercal, N., Aykin-Burns, N., Gurer-Orhan, H., McDonald, J.D., 2002. Oxidative stress in a phenylketonuria animal model. Free Radic Biol Med 32, 906–911. 10.1016/s0891-5849(02)00781-5

Humbert, M., Morán, M., de la Cruz-Ojeda, P., Muntané, J., Wiedmer, T., Apostolova, N., McKenna, S.L., Velasco, G., Balduini, W., Eckhart, L., Janji, B., Sampaio-Marques, B., Ludovico, P., Žerovnik, E., Langer, R., Perren, A., Engedal, N., Tschan, M.P., 2020. Assessing Autophagy in Archived Tissue or How to Capture Autophagic Flux from a Tissue Snapshot. Biology (Basel) 9, 59. 10.3390/biology9030059

Kuzu, O.F., Granerud, L.J.T., Saatcioglu, F., 2025. Navigating the landscape of protein folding and proteostasis: from molecular chaperones to therapeutic innovations. Signal Transduct Target Ther 10, 358. 10.1038/s41392-025-02439-w

Livak, K.J., Schmittgen, T.D., 2001. Analysis of relative gene expression data using real-time quantitative PCR and the 2(-Delta Delta C(T)) Method. Methods 25, 402–408. 10.1006/meth.2001.1262

Lu, L.-H., Xia, Z.-X., Guo, J.-L., Xiao, L.-L., Zhang, Y.-J., 2020. Metabolomics analysis reveals perturbations of cerebrocortical metabolic pathways in the Pahenu2 mouse model of phenylketonuria. CNS Neurosci Ther 26, 486–493. 10.1111/cns.13214

Masini, S., Bruschi, M., Menotta, M., Canonico, B., Montanari, M., Ligi, D., Monittola, F., Mannello, F., Piersanti, G., Crinelli, R., Magnani, M., Fraternale, A., 2025. Redox modulation by a synthetic thiol compound reduces LPS-induced pro-inflammatory cytokine expression in macrophages via AP-1/NLRP3 axis and influences the crosstalk with endothelial cells. Free Radic Res 1–19. 10.1080/10715762.2025.2529914

McDonald, J.D., Charlton, C.K., 1997. Characterization of mutations at the mouse phenylalanine hydroxylase locus. Genomics 39, 402–405. 10.1006/geno.1996.4508

Monittola, F., Masini, S., Montanari, M., Nasoni, M.G., Bianchi, M., De Matteis, R., Ricci, A., Ligi, D., Luchetti, F., Canonico, B., Magnani, M., Menotta, M., Fraternale, A., Crinelli, R., 2025. Immunoproteasome remodeling in senescing human macrophages reveals the loss of PA28αβ capping as a hallmark of immunosenescence. Commun Biol 8, 1371. 10.1038/s42003-025-08765-7

Ortega, M.A., Fraile-Martinez, O., de Leon-Oliva, D., Boaru, D.L., Lopez-Gonzalez, L., García-Montero, C., Alvarez-Mon, M.A., Guijarro, L.G., Torres-Carranza, D., Saez, M.A., Diaz-Pedrero, R., Albillos, A., Alvarez-Mon, M., 2024. Autophagy in Its (Proper) Context: Molecular Basis, Biological Relevance, Pharmacological Modulation, and Lifestyle Medicine. Int J Biol Sci 20, 2532–2554. 10.7150/ijbs.95122

Owegie, O.C., Kennedy, Q.P., Davizon-Castillo, P., Yang, M., 2025. Thiol Isomerases: Enzymatic Mechanisms, Models of Oxidation, and Antagonism by Galloylated Polyphenols. Antioxidants (Basel) 14, 1193. 10.3390/antiox14101193

Petrova, B., Warren, A., Vital, N.Y., Culhane, A.J., Maynard, A.G., Wong, A., Kanarek, N., 2021. Redox Metabolism Measurement in Mammalian Cells and Tissues by LC-MS. Metabolites 11, 313. 10.3390/metabo11050313

Rocha, J.C., Martins, M.J., 2012. Oxidative stress in phenylketonuria: future directions. J Inherit Metab Dis 35, 381–398. 10.1007/s10545-011-9417-2

Roelofs, J., Suppahia, A., Waite, K., Park, S., 2018. Native Gel Approaches in Studying Proteasome Assembly and Chaperones: Methods and Protocols, in: Methods in Molecular Biology (Clifton, N.J.). pp. 237–260. 10.1007/978-1-4939-8706-1_16

Ross, A.B., Langer, J.D., Jovanovic, M., 2020. Proteome Turnover in the Spotlight: Approaches, Applications, and Perspectives. Mol Cell Proteomics 20, 100016. 10.1074/mcp.R120.002190

Schlegel, G., Scholz, R., Ullrich, K., Santer, R., Rune, G.M., 2016. Phenylketonuria: Direct and indirect effects of phenylalanine. Exp Neurol 281, 28–36. 10.1016/j.expneurol.2016.04.013

Seco-Cervera, M., González-Cabo, P., Pallardó, F.V., Romá-Mateo, C., García-Giménez, J.L., 2020. Thioredoxin and Glutaredoxin Systems as Potential Targets for the Development of New Treatments in Friedreich’s Ataxia. Antioxidants (Basel) 9, 1257. 10.3390/antiox9121257

Sirtori, L.R., Dutra-Filho, C.S., Fitarelli, D., Sitta, A., Haeser, A., Barschak, A.G., Wajner, M., Coelho, D.M., Llesuy, S., Belló-Klein, A., Giugliani, R., Deon, M., Vargas, C.R., 2005. Oxidative stress in patients with phenylketonuria. Biochim Biophys Acta 1740, 68–73. 10.1016/j.bbadis.2005.02.005

Sitta, A., Barschak, A.G., Deon, M., Terroso, T., Pires, R., Giugliani, R., Dutra-Filho, C.S., Wajner, M., Vargas, C.R., 2006. Investigation of oxidative stress parameters in treated phenylketonuric patients. Metab Brain Dis 21, 287–296. 10.1007/s11011-006-9035-0

Tanaka, K., 2009. The proteasome: overview of structure and functions. Proc Jpn Acad Ser B Phys Biol Sci 85, 12–36. 10.2183/pjab.85.12

Thau-Zuchman, O., Pallier, P.N., Savelkoul, P.J.M., Kuipers, A.A.M., Verkuyl, J.M., Michael-Titus, A.T., 2022. High phenylalanine concentrations induce demyelination and microglial activation in mouse cerebellar organotypic slices. Front Neurosci 16, 926023. 10.3389/fnins.2022.926023

Tutas, J., Tolve, M., Özer-Yildiz, E., Ickert, L., Klein, I., Silverman, Q., Liebsch, F., Dethloff, F., Giavalisco, P., Endepols, H., Georgomanolis, T., Neumaier, B., Drzezga, A., Schwarz, G., Thorens, B., Gatto, G., Frezza, C., Kononenko, N.L., 2025. Autophagy regulator ATG5 preserves cerebellar function by safeguarding its glycolytic activity. Nat Metab 7, 297–320. 10.1038/s42255-024-01196-4

Yang, H.-C., Stern, A., Chiu, D.T.-Y., 2021. G6PD: A hub for metabolic reprogramming and redox signaling in cancer. Biomed J 44, 285–292. 10.1016/j.bj.2020.08.001

Yazgili, A.S., Meul, T., Welk, V., Semren, N., Kammerl, I.E., Meiners, S., 2021. In-gel proteasome assay to determine the activity, amount, and composition of proteasome complexes from mammalian cells or tissues. STAR Protoc 2, 100526. 10.1016/j.xpro.2021.100526

