## Supplementary files for "Integrated metabolic and proteostatic profiling reveals remodeling of proteolytic pathways associated with redox-bioenergetic dysfunction in a PAH^enu2^ mouse model of phenylketonuria"

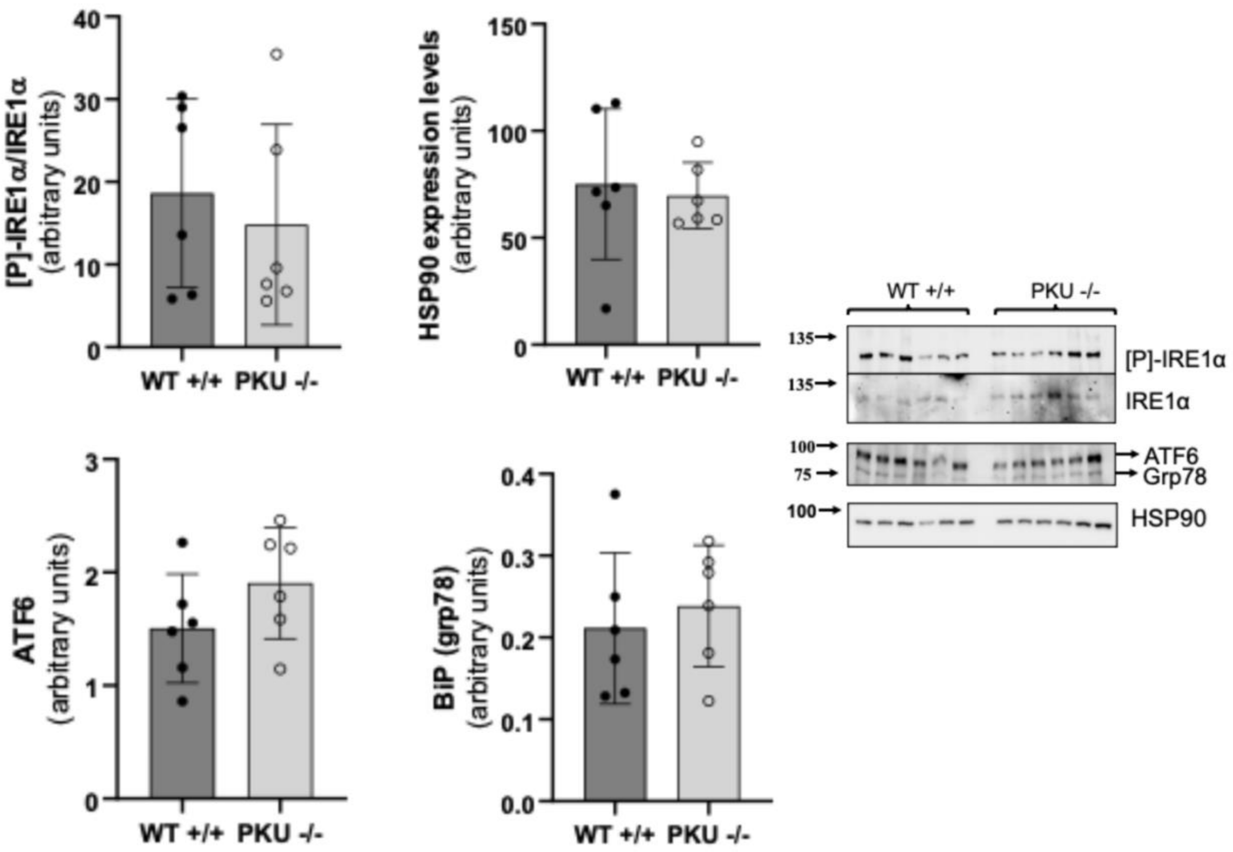


**Figure S1. Analysis of unfolded protein response markers in WT and PKU mouse brains.**
Western blot analysis of phosphorylated IRE1α and total IRE1α expression in WT and PKU brain samples was performed by loading 25 µg of total protein lysate onto a 6% (w/v) acrylamide gel. The phosphorylation level of IRE1α (active form) was normalized to total IRE1α protein levels following membrane stripping and reprobing. Western blot analysis of HSP90, ATF6, and BiP/GRP78 expression was performed by loading 20 µg of total protein lysate onto a 10% (w/v) acrylamide gel. β-actin was used as the loading control for normalization. Results are presented in the graph as mean ± SD.

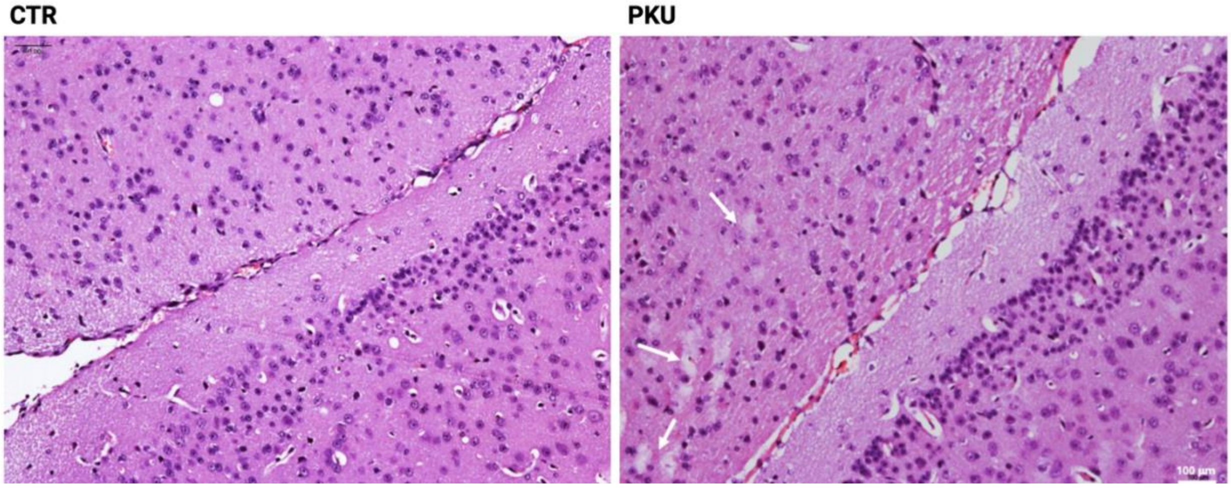


**Figure S2. Representative histological sections of the corresponding brain region in control and PKU specimens.** Neuronal rarefaction is observed in the PKU tissue (white arrows). H&E staining. Scale bar: 100 µm.

.

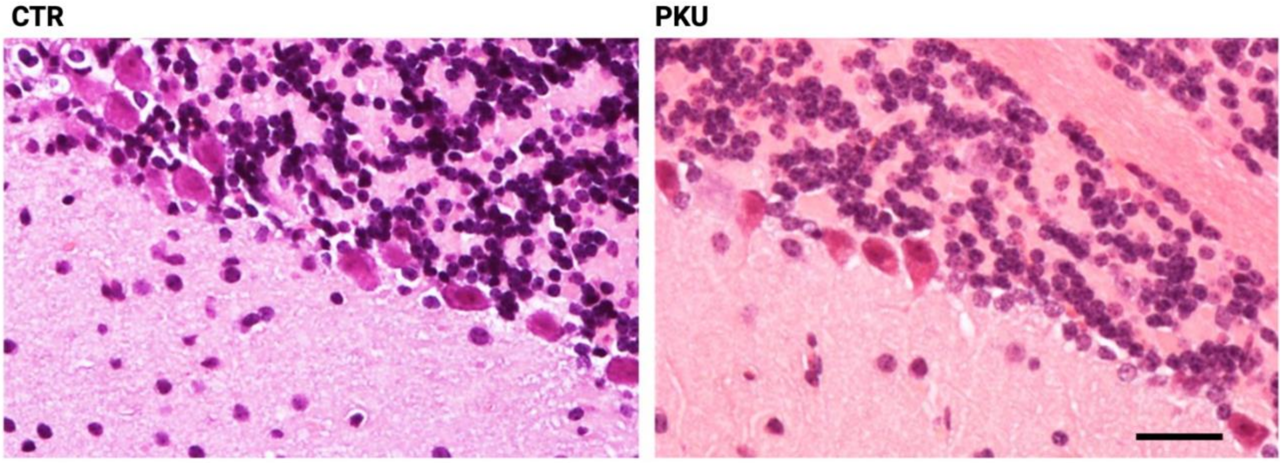


**Figure S3. Representative images of cerebellar sections from control and PKU mice.** A reduced number of Purkinje cells is observed in the Purkinje cell layer of PKU mice compared to control. H&E staining. Scale bar: 100 μm.

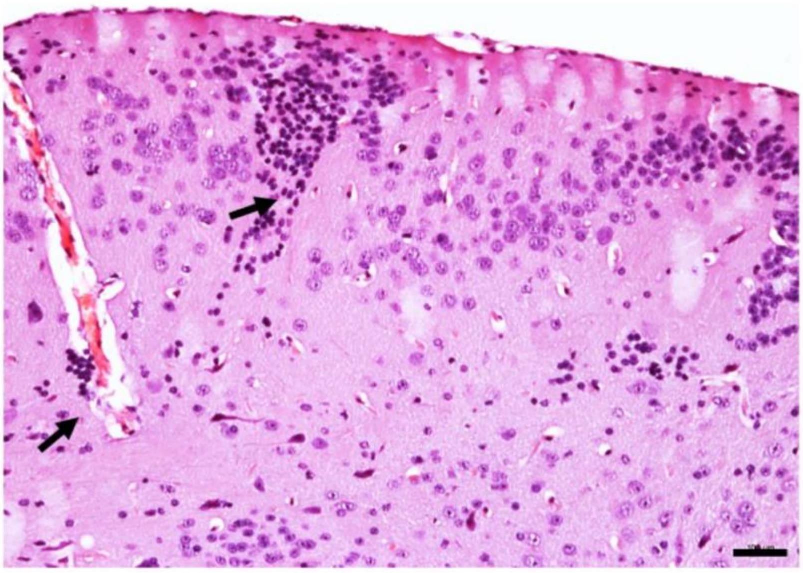

**Figure S4. Perivascular lymphocytic infiltrate in the PKU mouse brain.** Perivascular and parenchymal inflammatory infiltrates are observed in the cerebral cortex of PKU mice (black arrows). H&E staining. Scale bar: 100 µm.

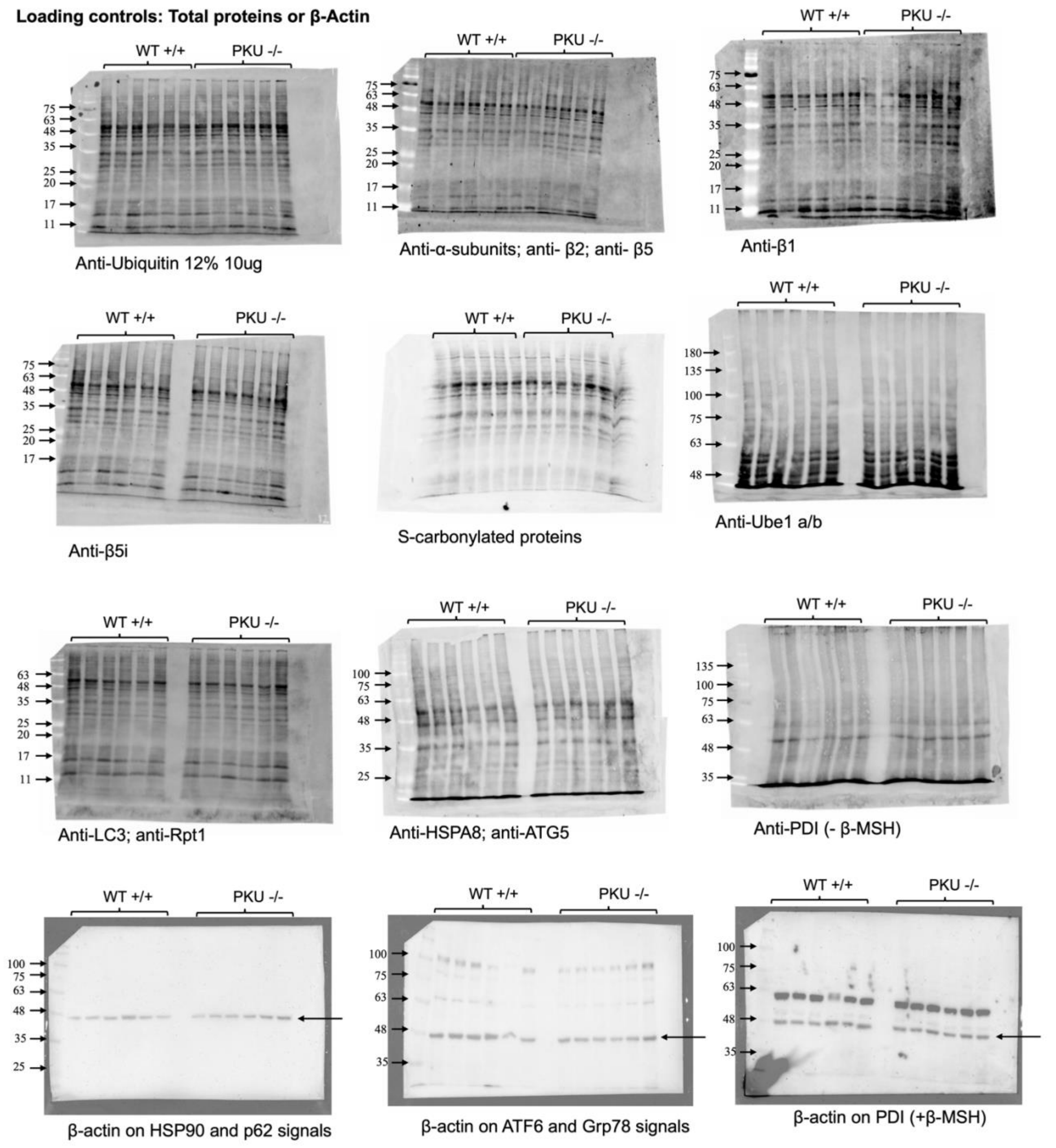


**Figure S5. Raw western blot data**

Representative images of total protein staining obtained with the No-Stain kit and β-actin immunodetection, used as loading controls for western blot normalization.

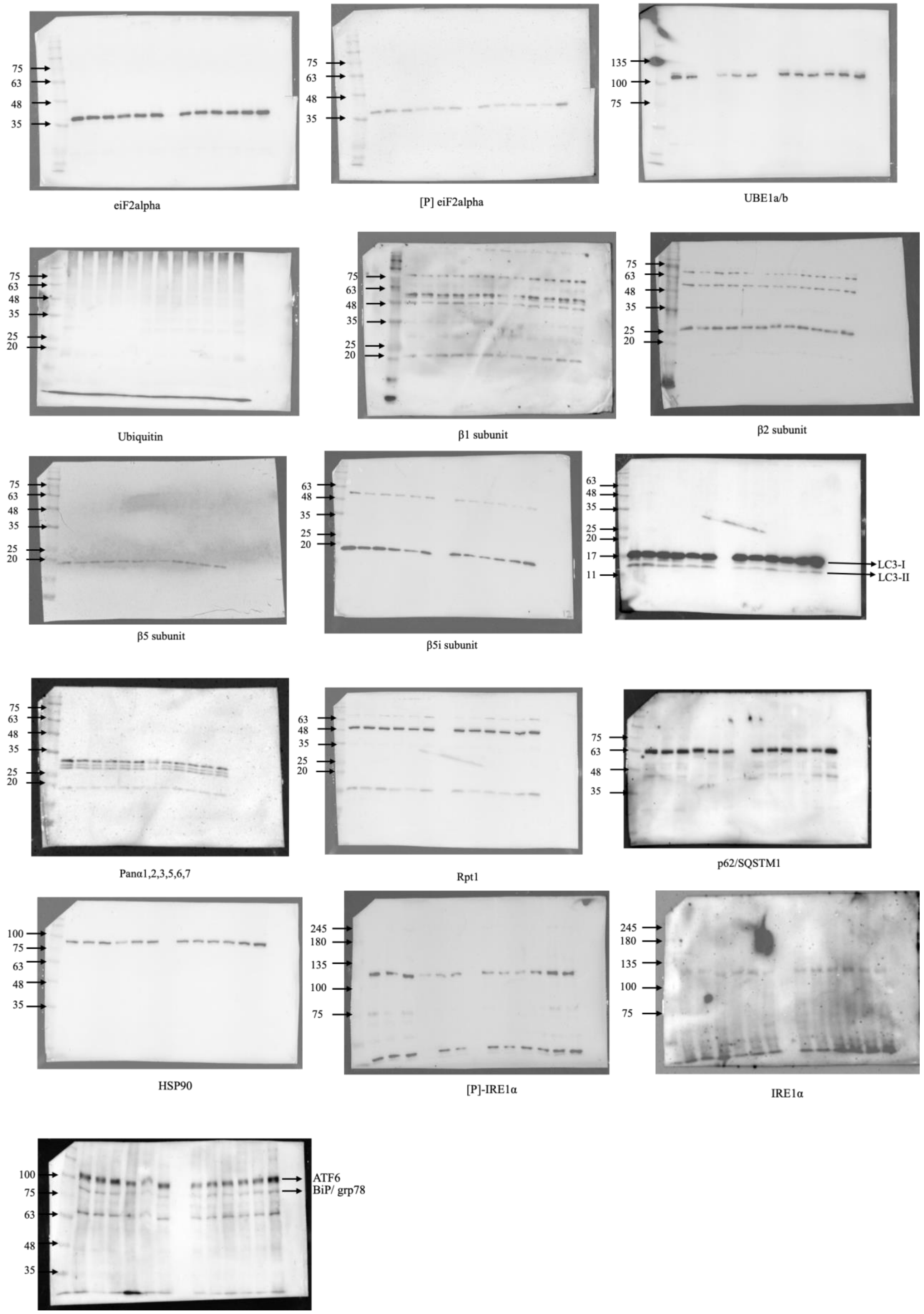


**Figure S6. Uncropped images of Western blots.**
